# Automated and unbiased classification of motor neuron phenotypes with single cell resolution in ALS tissue

**DOI:** 10.1101/2020.08.17.253773

**Authors:** Cathleen Hagemann, Giulia E. Tyzack, Doaa M. Taha, Helen Devine, Linda Greensmith, Jia Newcombe, Rickie Patani, Andrea Serio, Raphaëlle Luisier

## Abstract

Histopathological analysis of tissue sections is an invaluable resource in neurodegeneration research. Importantly, cell-to-cell variation in both the presence and severity of a given phenotype is however a key limitation of this approach, reducing the signal to noise ratio and leaving unresolved the potential of single-cell scoring for a given disease attribute. Here, we developed an image processing pipeline for automated identification and profiling of motor neurons (MNs) in amyotrophic lateral sclerosis (ALS) pathological tissue sections. This approach enabled unbiased analysis of hundreds of cells, from which hundreds of features were readily extracted. Next by testing different machine learning methods, we automated the identification of phenotypically distinct MN subpopulations in VCP- and SOD1-mutant transgenic mice, revealing common aberrant phenotypes in cellular shape. Additionally we established scoring metrics to rank cells and tissue samples for both disease probability and severity. Finally, by adapting this methodology to human post-mortem tissue analysis, we validated our core finding that morphological descriptors strongly discriminate ALS from control healthy tissue at the single cell level. In summary, we show that combining automated image processing with machine learning methods substantially improves the speed and reliability of identifying phenotypically diverse MN populations. Determining disease presence, severity and unbiased phenotypes at single cell resolution might prove transformational in our understanding of ALS and neurodegenerative diseases more broadly.

## INTRODUCTION

Amyotrophic lateral sclerosis (ALS) is a relentlessly progressive neurodegenerative disease, which remains incurable due to an incomplete understanding of the underlying molecular pathogenesis. Using the VCP mutant mouse transgenic model of ALS and human post-mortem tissue from sporadic ALS (sALS) patients, we previously showed that spinal motor neurons (MNs) exhibit reduced nuclear to cytoplasmic (N/C) ratios of FUS (Fused in Sarcoma) and SFPQ (Splicing factor Proline and Glutamine rich) proteins. Furthermore, we showed that SFPQ is also mislocalized in SOD1-mutant ALS models, while FUS is not (Luisier *et al*., 2018; Tyzack *et al*., 2019).

Although these studies revealed novel molecular hallmarks of ALS, they also present important limitations shared by the majority of such traditional neuropathological studies. Firstly, key observations are based on manual image processing, which is time-consuming (therefore not scalable to large studies) and also introduces potential operator-dependent variability. Secondly, although hundreds of measurements can be extracted from such rich images, previous studies have typically focused on the single measurement deemed to be the most relevant. Furthermore, such analyses are usually based on averaging the signal(s) among all cells originating from identical animals or individuals. However, the degree to which individual MNs follow population-averaged trends in protein mislocalization remains unresolved. Indeed important insights into biological processes such as cellular differentiation and disease progression originate from the characterisation of heterogeneous cell populations coexisting within the same condition (Rubin, 1990; Chang *et al*., 2008; Slack *et al*., 2008, Loo *et al*., 2009*a*; Snijder and Pelkmans, 2011; Ho *et al*., 2020). These studies have been very effective when applied to assays of cultured cells and for cancer subtype classification, however there is not, to our knowledge, an analogous study on tissue sections from the nervous system (Yamamoto *et al*., 2017; Ščupáková *et al*., 2020). Finally these studies do not provide disease scoring at the cellular or even individual or animal level, which would be advantageous for understanding single cell phenotypic heterogeneity within a disease and, in the longer term, for developing superior patient-specific diagnostic criteria.

Recent developments using deep learning methods have enabled segmentation and object detection from complex imaging data-sets, with exciting prospects within ALS translational research (Berg *et al*., 2019*b*; Von Chamier *et al*., 2020). Here, by expanding on these methods, we developed a pipeline for image processing of MNs from spinal cord sections, greatly improving efficiency, whilst providing an unbiased approach to data acquisition. We then extracted hundreds of cell measurements and compared different machine learning methods to automatically identify MNs subpopulations from high-content imaging data. Using this approach we reveal that VCP- and SOD1-mutant MNs share phenotypes captured by several related morphological descriptors including Zernike moments, and we validate this finding in human ALS post-mortem tissues. By providing our fluorescence microscopy raw images together with open-source implementations of the methods, we aim to allow others to readily apply these methods in other biological contexts in order to increase the analytical power of tissue section analysis. We propose that such unbiased approaches may substantially deepen our understanding of ALS and other diseases in which we routinely use histopathological analysis.

## RESULTS

### Automated phenotyping of spinal cord MNs from transgenic ALS mouse models

Large variability in phenotype between transgenic ALS mouse models is well described in the literature (De Giorgio *et al*., 2019). We previously showed that VCP- and SOD1-mutant ALS mouse models exhibit distinct pathological phenotypes in terms of FUS and/or SFPQ mislocalization (Luisier *et al*., 2018; Tyzack *et al*., 2019). In these studies, we analysed spinal cord sections from control, SOD1- and VCP-mutant mouse models immuno-labeled for FUS and SFPQ, where nuclear and cytoplasmic compartments were manually identified with DAPI and ChAT (Luisier *et al*., 2018; Tyzack *et al*., 2019). Using these same images (**Table S1**), we first aimed to test whether spinal cord MNs from these ALS mouse models exhibit common cellular phenotypes which were not captured by our previous analyses. In order to achieve this, we moved beyond single-protein localisation analysis at the population average level to comprehensive single cell analyses by developing a pipeline for automated image segmentation and morphological profiling combining the open-source Ilastik and CellProfiler softwares (Carpenter *et al*., 2006, Berg *et al*., 2019*a*) (**Fig. 1A** and **Supplementary Fig. 1A**). This allowed for the rapid, automated identification of distinct cellular compartments, and extraction of 750 morphological features for every MN identified (n=121), thereby greatly improving processing speed and throughput compared to manual processing. Features included fluorescent intensity data and signal distribution (for FUS, SFPQ, ChAT and DAPI), shape and morphometric descriptors (size, shape, perimeter and texture of subcellular structures including the nucleus and cytoplasm).

**Figure 1.**
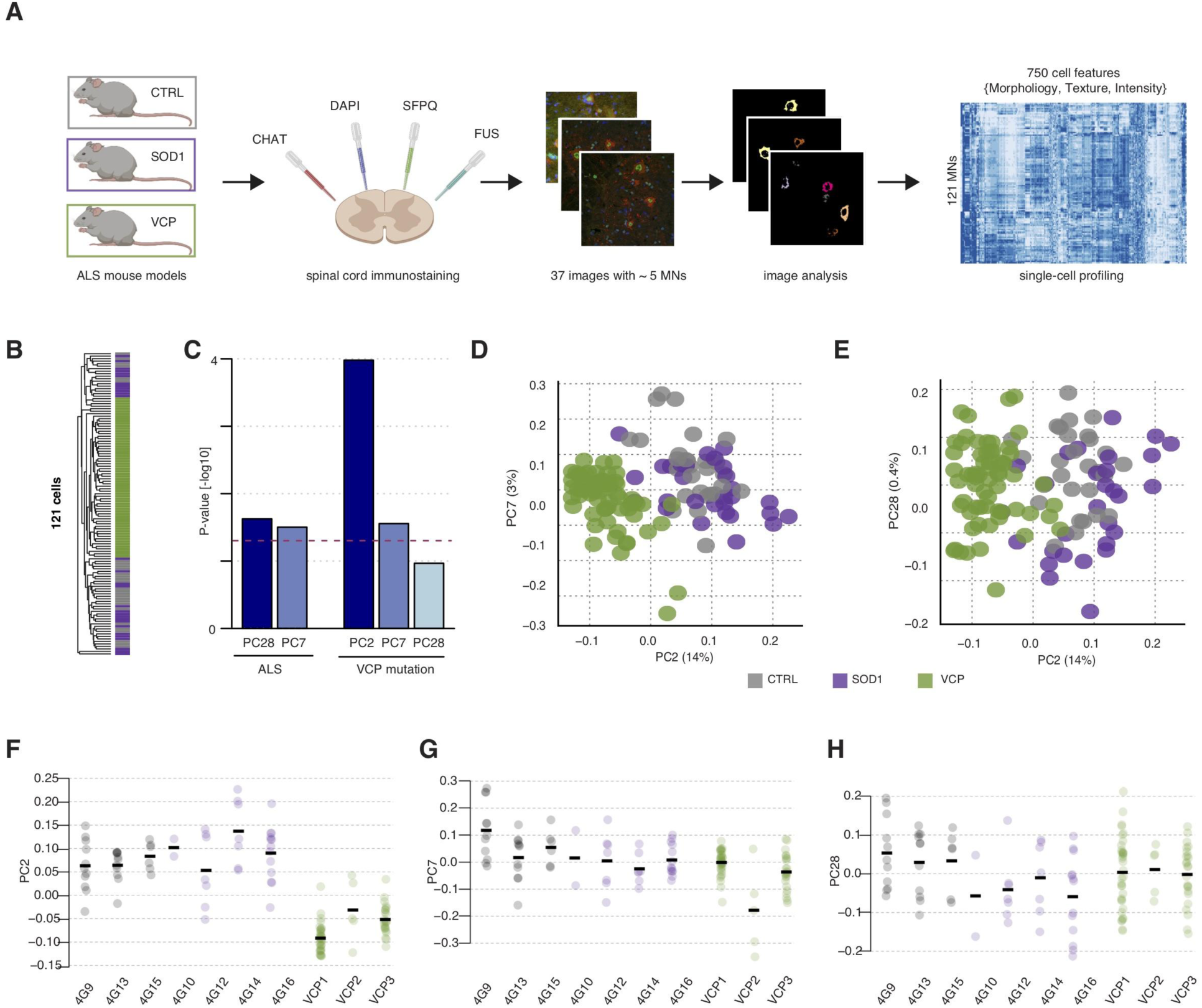
High-content MNs profiling captures cell features that discriminate the ALS phenotype in mice spinal cord sections. (**A**) Workflow for generating morphological profiles of MNs in ALS pathological spinal cord sections. Spinal cord sections from SOD1*^G93A^* mutant (n = 4 mice), mutant human VCP*^A232E^* (n = 3 mice), and wild-type (n = 3 control mice) mice were co-stained against ChAT, DAPI, SFPQ and FUS and then imaged. A combination of *Ilastik* (Berg *et al*., 2019*a*) and *CellProfiler* (Carpenter *et al*., 2006) enables automatic identification and feature extraction from cytoplasmic and nuclear compartments. (**B**) Unsupervised hierarchical clustering of 121 cells using 750 scaled measurements shows evidence for VCP-mutation dominant signal over SOD1-mutation in ALS MNs populations. Grey bars = control MNs; magenta bars = SOD1 MNs; green bars = VCP MNs. Euclidean distance and average clustering method. (**C**) Barplots showing the association between principal components and either common ALS (comALS) or VCP (vcpALS) phenotype. Linear mixed effects analysis of the relationship between comALS or vcpALS phenotypes and each of the 31 first principal components accounting for idiosyncratic variations due to animals shows significant association of PC7 and PC28 with comALS phenotype and PC2 only with vcpALS phenotype. (**D, E**) SVD performed on 750 standardized measurements across 121 cells. Cells are plotted by their coordinates along PC2 (14% of variance), PC7 (3% of variance), and PC28 (0.4% of variance). Colors of data points indicate similar mouse genetic background: control (grey), SOD1 mutant mice (magenta), and VCP mutant mice (green). (**F-H**) Individual cell coordinates along PC2, PC7 and PC8 grouped by animal and colored according to genetic background.

We next examined whether phenotypic states either associated with ALS-mutant cells or control cells are accompanied by reproducible changes in morphological descriptors. Unsupervised hierarchical clustering (Euclidean distance and average clustering) of the 121 MNs using the 750 measured morphological features segregated cells based on the presence or absence of the VCP-mutation rather than segregating SOD1- and VCP-mutant cells together (**Fig. 1B**). This suggests that the ALS phenotype associated with the VCP mutation, hereafter referred to as *vcpALS* phenotype, dominates a common ALS phenotype shared between SOD1 and VCP mutant MNs, hereafter referred to as *comALS* phenotype. Hierarchical clustering is, however, often dominated by a single morphological profile and thus can fail to reveal subtle but relevant features in the data (Ronan *et al*., 2016). In contrast to this, singular value decomposition (SVD) permits deconvolution and ranking of orthogonal morphological profiles (Luisier *et al*., 2014). Here we find that the information contained in the data is well distributed among the 121 components derived from SVD analysis (Shannon Entropy=0.649), and that the first 31 components capture 90% of the variance in the data (**Supplementary Fig. 2**). Using linear mixed modelling we find significant association between principal component (PC) 7 (3% of variance) and PC28 (0.4% of variance) with *comALS* phenotype (SOD1 and VCP mutation), while PC2 (14% of variance) is strictly associated with *vcpALS* phenotype (**Fig. 1C**). While this analysis confirms the dominance of morphological changes associated with the VCP mutation, as further visualised in the scatter plots (**Figs. 1D,E** and **Fig. 1F**), it indicates that SOD1- and VCP-mutant cells together exhibit subtle and consistent morphological characteristics likely to reflect common ALS attributes (**Figs. 1G,H**).

### Machine learning methods can identify common phenotypes between SOD1 and VCP mutant ALS mouse models

Having established that high-content MNs profiling data from fluorescence microscopy can capture subtle but consistent phenotypes across VCP and SOD1 genetic backgrounds, we next aimed to train and compare different machine learning methods to automatically identify MNs exhibiting this *comALS* phenotype in spinal cord sections. Visual inspection of the distribution of the principal components associated with either *comALS* (PC7 and PC28) or *vcpALS* (PC2) phenotype revealed that PC2 and PC28, but not PC7, exhibit bimodal distributions (**Figs. 2A-C**). This observation suggests the presence of at least two distinct subpopulations among the 121 cells. Here we propose that MNs populations can be described in terms of a mixture of two phenotypically distinct subpopulations, namely “healthy” and “sick (diseased)” cells (hereafter referred to as **disMNs**). Given that healthy cells can exist in spinal cord sections of VCP/SOD1 mutant mice, we selected the probabilistic label-independent Gaussian Mixture Model (GMM) classifier, and compared it with two label-dependent classifiers, namely Logistic Regression (LR) and Multi-Layer Perceptron (MLP). We tested GMM with the following combinations of principal components selected based on their association with either *comALS* or *vcpALS* phenotype: *GMM-PC2*, *GMM-PC2,PC7*, *GMM-PC7,PC28*, and *GMM-PC28*.

**Figure 2.**
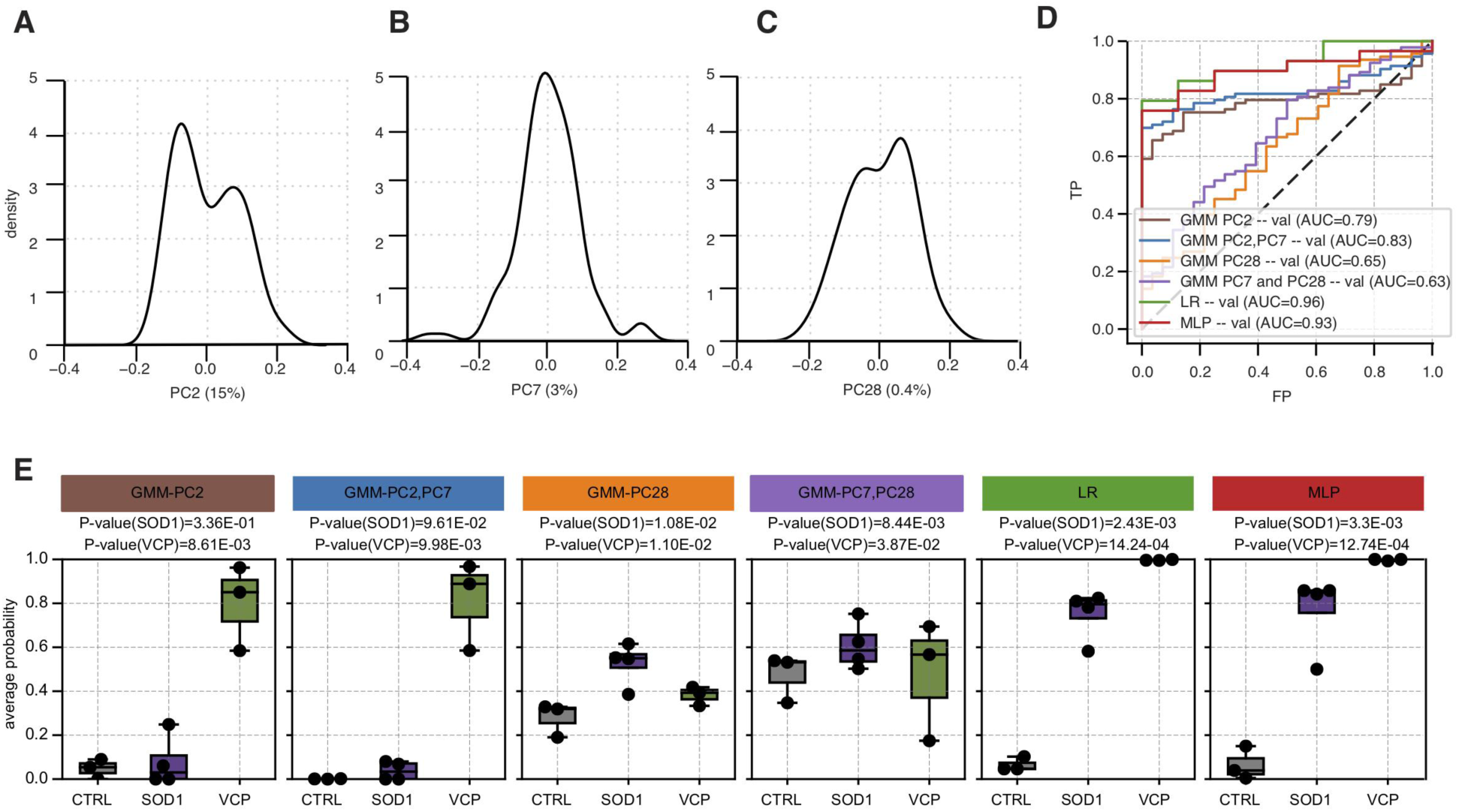
Machine-learning based methods enable automated discrimination of VCP- and SOD1-mutant groups from control groups. (**A,B,C**) Density distributions of MNs contributions on PC2 (15% of variance), PC7 (3% of variance) and PC28 (0.4% of variance) respectively. (**D**) Performance analysis of each classifier in their ability to discriminate ALS-mutant cells from control cells using receiver operating characteristic (ROC) curves and areas under the curves (AUC). (**E**) The ability of each clustering algorithm to detect SOD-1 or VCP-mutation effect is assessed by comparing the disease probabilities of either the SOD1- or VCP-mutant groups with those of the control group. The disease probability of each animal is obtained by using the mean probabilities of its cells to be sick according to 1) *GMM-PC2,PC7*, 2) *GMM-PC28*, 3) Logistic regression or 4) Multilayer perceptron classifiers. Data shown as box plots in which the centre line is the median, limits are the interquartile range and whiskers are the minimum and maximum. Dots are the animal disease profile. *P*-values obtained from Welch’s t test.

When considering accuracy in the ability of the classifiers to discriminate VCP- and SOD1-mutant cells from control cells, LR and MLP performed equally well (AUC*_LR_*=0.96 and AUC*_MLP_*=0.93), while GMM-derived methods resulted in lower scores overall (AUC*_GMM-PC2,PC7_*=0.83 > AUC*_GMM-PC2_*=0.79 > AUC*_GMM-PC28_*=0.65> AUC*_GMM-PC7,PC28_*=0.63; **Fig. 2B**). AUC (Area Under the Curve) and other related scores are widely used to rank classifiers, however these need to be treated with caution in the current context given the existence of healthy cells in ALS-derived tissue sections (and indeed theoretical disMNs in control mice). We therefore assigned each animal a probability to be sick using the mean probability of their respective cells, as provided by each classifier, and tested whether the VCP and/or SOD1-mutant group exhibited significantly higher probability to be sick compared to the control group. Comparison of the classifiers’ ability to discriminate control from ALS-mutant animals using disease probability is expected to better reflect the biological significance of these classifiers. Using this method we showed that only the VCP-mutant group exhibited significantly higher disease probability than the control group when probabilities were derived from either *GMM-PC2* or *GMM-PC2,PC7* (*P*_GMM-PC2|*VCP*_ =8.61e-03, *P*_GMM-PC2|*SOD1*_ =3.4e-01, *P*_GMM-PC2,PC7|VCP_=9.9e-03, and *P*_GMM-PC2,PC7|SOD1_=9.6e-02; **Fig. 2E**). Furthermore, *GMM-PC28* resulted in a significant difference between control and SOD1-mutant groups, and to some extent between VCP-mutant and control groups (*P*_GMM-PC28|VCP_ =1.1e-02, *P*_GMM-PC28|SOD1_ =1.0e-02). In summary, only *GMM-PC28* enabled recovery of *comALS* effect, however at higher significance for SOD1 group. In contrast, both LR and MLP classifiers resulted in large significant differences between control and ALS mutant groups irrespective of their genetic backgrounds (*P*_LR|VCP_ =1.4e-04, *P*_ML|VCP_=1.2e-04, *P*_LR|SOD1_ =2.4e-03 and *P*_MLP|SOD1_=3.3e-03). These results suggest that PC2 and PC7 capture a cellular phenotype characteristic of *vcpALS*. Furthermore, MLP, LR and to some extent *GMM-PC28* are able to capture a unifying phenotype amongst VCP and SOD1 mice, i.e. *comALS*. In the following sections we focus on the three *comALS* classifiers, namely LR, MLP, *GMM-PC28,* and compare these with the *vcpALS* classifier *GMM-PC2,PC7* to study their underlying differences.

### LR classifier best captures the broader ALS phenotype at single-cell level

To describe the disease status of an animal or a tissue section, it is important to acknowledge that two cells (or indeed tissues/animals and patients) with similar disease probability can also exhibit different degrees of aberrant phenotype(s). Here we associated each MN with the following two metrics together forming a per-cell ‘disease profile’: 1) the disease probability *P* given the observed phenotype as outputted by the classifiers; 2) a ‘severity’ score *S* directly derived from *P* and expected to reflect on the possibility for a population of cells with identical *P* to exhibit different degrees of disease phenotypes (see Materials and Methods, and **Supplementary Fig. 3**).

We next performed hierarchical clustering (Euclidean distance and average clustering) of the MNs using the 121 disease profiles to test the ability of each classifier to group cells into disease versus healthy groups. We found that the *vcpALS* classifier *GMM-PC2,PC7* segregated cells in two groups according to the VCP genetic background (**Fig. 3A**). As expected, some MNs derived from VCP-mutant mice cluster with cells derived from either control or SOD1-mutant group; these are potentially healthy cells present in mutant mice reflecting the cellular heterogeneity in tissue. *GMM-PC28* classifier segregated MNs in two genetically heterogeneous groups, the smaller group being exclusively composed of cells derived from ALS-mutant mice (4 VCP-mutant cells and 2 SOD1 mutant cells), and therefore called *comALS* group. LR and MLP similarly clusters cells in three large groups, one group almost exclusively composed of cells derived from control mice (and some SOD1-mutant cells), called the healthy group, one group almost exclusively composed of VCP-mutant cells, called the vcpALS group, and the third group being a mixture of SOD1- and VCP-mutant cells, i.e the comALS group. This analysis confirms that LR, MLP, and to some extent *GMM-PC28*, capture the *comALS* phenotype. This is distinct from the *vcpALS* classifier *GMM-PC2,PC7* that discriminates VCP-mutant cells from the control and SOD1-mutant cells. Notably, using LR we can clearly separate the control group from the vcpALS/comALS groups with the highest degree of confidence (i.e. separation between the groups), whilst MLP groups cells in three clusters, healthy, vcpALS and comALS. This indicates that the MLP classifier first captures vcpALS phenotype whilst LR, and to some extent *GMM-PC28*, first capture the comALS phenotype.

**Figure 3.**
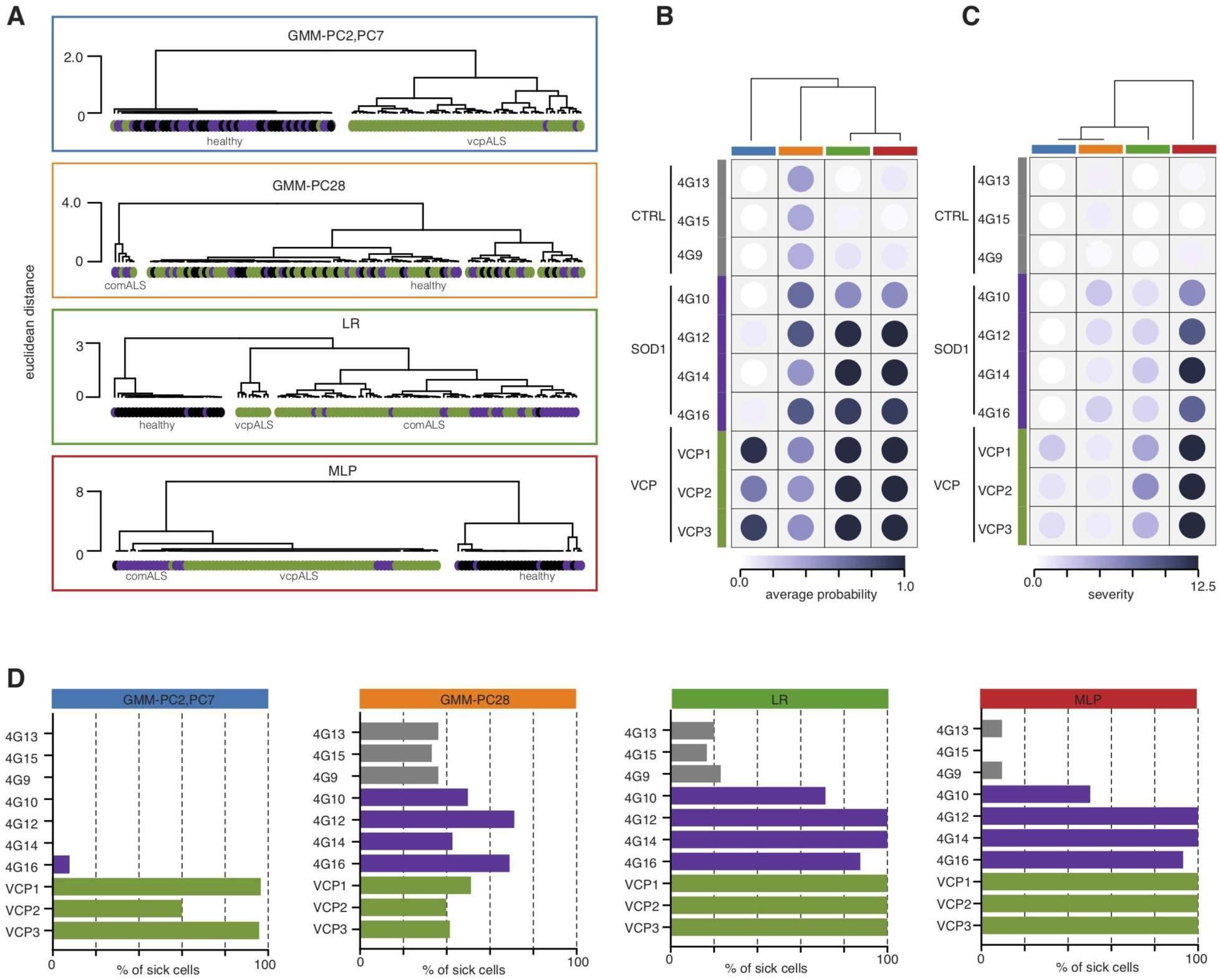
Comparisons of the different classifiers for predicted percentage of sick cells, disease probability, and disease severity. (**A**) Comparison of the disease profile similarity among the ALS-mutant cells versus control cells as obtained from *GMM-PC2-PC7* (blue rectangle), *GMM-PC28* (orange rectangle), LR (green rectangle) or MLP (red rectangle) classifiers using unsupervised hierarchical clustering of the 121 cells. Grey circles = control MNs; magenta circles = SOD1-mutant MNs; green circles = VCP-mutant MNs. (**B**, **C**) Heatmap of the animal-level disease probabilities and disease severity scores as predicted by *GMM-PC2-PC7* (blue bar), *GMM-PC28* (orange bar), LR (green bar) or MLP (red bar) classifiers. The disease probability and disease scores of each animal are obtained by using the mean probabilities to be sick and severity scores respectively of its cells. The different classifiers are hierarchically clustered using the average algorithm on the Euclidean distance between the disease probabilities or disease severity scores across the 10 animals. (**D**) Bar plots displaying the animal-level percentage of sick cells as predicted by *GMM-PC2,PC7*, *GMM-PC28*, LR or MLP classifiers. Grey bars = control mice. Magenta bars = SOD1-mutant mice. Green bars = VCP-mutant mice.

From the two per-cell disease score metrics, we next derived for each animal 1) the animal probability to be sick, 2) the animal disease severity, and 3) the animal percentage of disMNs, and used them to quantify the similarity between the four classifiers *GMM-PC2,PC7*, *GMM-PC28*, LR, and MLP. Using hierarchical clustering based on the per-animal probability to be sick further confirmed the largest differences between the vcpALS classifier *GMM-PC2,PC7* and the three comALS classifiers *GMM-PC28,* LR, MLP are similar (**Fig. 3B**). When the per-animal disease severity was used to compare the classifiers however, MLP was the most divergent classifier, exhibiting large severity for all ALS mutant mice irrespective of their genetic background (**Fig. 3B**). This indicates that although MLP predicts large differences in disease profiles between vcpALS and comALS groups of cells, when looking at the per-animal severity, both VCP- and SOD1-mutant animals exhibit large severity.

Finally we looked at the predicted percentage of disMNs obtained by each classifier and found that the vcpALS classifier *GMM-PC2,PC7* predicted disMNs only in VCP-mutant animals (**Fig. 3C**), while the three comALS classifiers (LR, MLP and *GMM-PC28)* predicted disMNs in ALS-mutant animals (**Fig. 3A**). Notably *GMM-PC28* predicted disMNs across all animals although with higher percentages in SOD1-mutant mice and VCP-mutant mice.

In summary, this analysis reveals that MLP, LR and to some extent GMM-PC28 deliver similar results in terms of their ability to discriminate ALS-mutant mice from control animals (similar per-animal scoring metrics). However, for individual cell scoring metrics, LR is the classifier which best captures measurements associated with comALS phenotype rather than dominant vcpALS phenotype, as evidenced by the large component driven by the VCP mutation in MLP classification.

### Morphometric descriptors are top classifiers for ALS phenotypes

The finding that LR and MLP best capture *comALS* cell phenotype prompted us to study what combinations of measurements actually carry the relevant information for ALS disease. We therefore looked into which measurements contribute most to the *vcpALS* versus *comALS* classifiers. In order to exclude the possibility that divergence in the top contributors between the *vcpALS* versus *comALS* classifiers could stem from the different mathematical foundations (GMM, LR and MLP), we trained two additional classifiers (one LR and one MLP hereafter called sLR and sMLP) with a subset of the data composed of control and VCP-mutant cells only, to specifically learn the *vcpALS* phenotype using these methods (**Supplementary Figs. 4A,B**). Hierarchical clustering of the classifiers using the predicted disease probability scores across the 121 MNs confirmed the similarity between *GMM-PC2,PC7*, sLR and sMLP in identifying *vcpALS* phenotype, compared with MLP, LR and *GMM-PC28* in identifying *comALS* phenotype (**Supplementary Fig. 4C**). Remarkably, although the *comALS* or *vcpALS* classifiers arise from diverse mathematical foundations, the relative contribution of the 750 measurements are very similar among these two groups of classifiers (**Fig. 4A**). This result indicates that the cell measurements which lead to the grouping in *comALS* or *vcpALS* groups are robust enough to be independent of the modeling approach.

**Figure 4.**
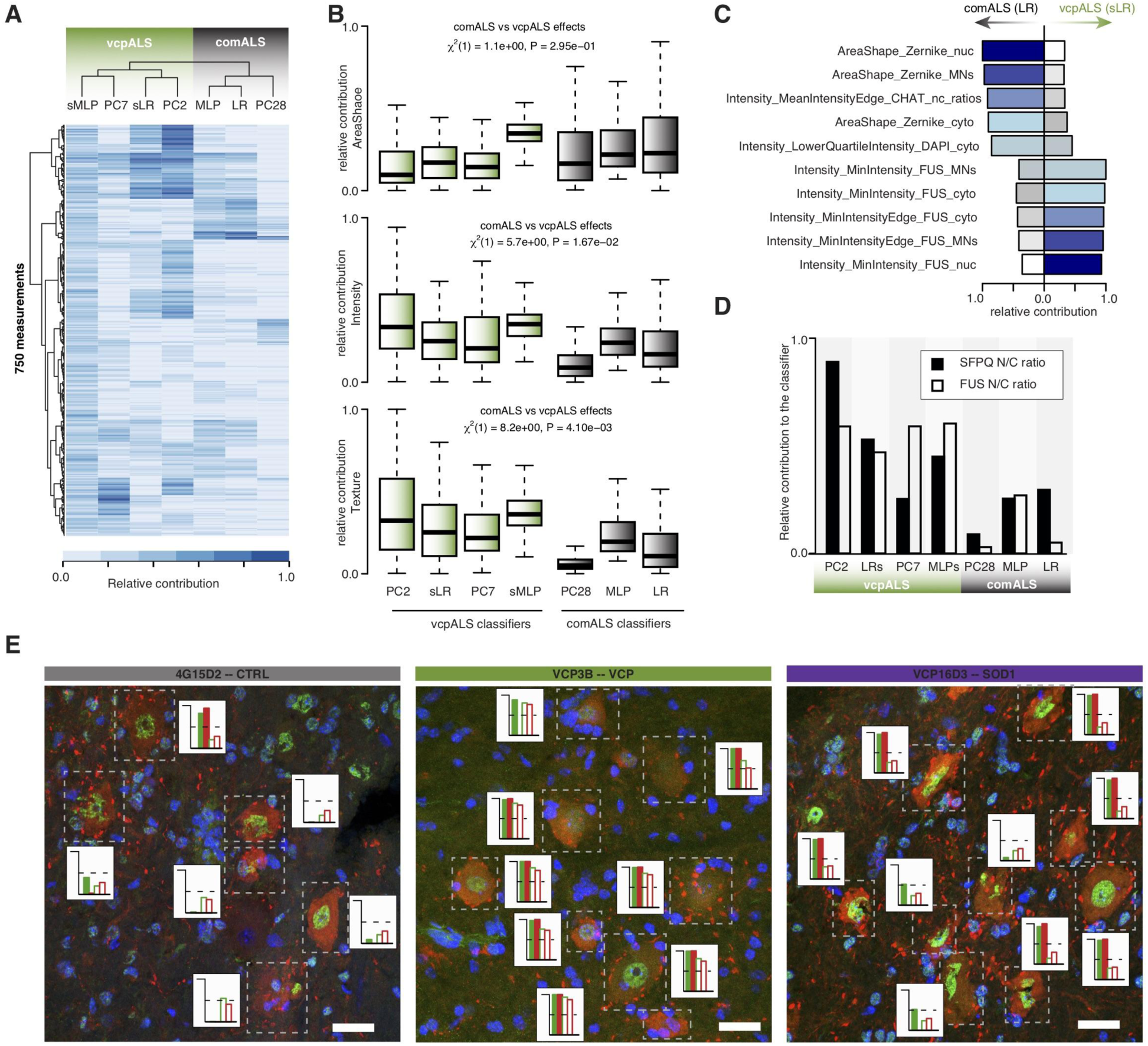
Area-shape related measurements contribute to the identification of ALS-mutant cells. (**A**) Heatmaps of the relative contribution of the 750 cell measurements of the different classifiers when applied or to the distinct principal components (PCs) to which GMM classifiers are fitted. The different classifiers/PCs are hierarchically clustered using the average algorithm of the euclidean distance between the relative contributions across the 750 measurements. vcpALS classifiers (green) discriminate VCP-mutant from SOD1-mutant and control cells. comALS classifiers (grey) discriminate ALS-mutant from control cells. (**B**) Boxplots showing the distributions of the relative contributions of area-shape (top), texture (middle) and intensity (bottom) related measurements in each classifier or PCs when applied. Linear mixed effects analysis of the relationship between comALS and vcpALS classifiers and the relative contribution of each measurement category to account for idiosyncratic variation due to classifiers. Data shown as box plots in which the centre line is the median, limits are the interquartile range and whiskers are the minimum and maximum. Green = VCP classifiers; grey = ALS classifiers. (**C**) Barplots showing the relative contribution of the top five measurements in LR (comALS classifier) and sLR (vcpALS classifiers). Zernike moments either in the nucleus or the whole MNs contribute largely to comALS but not vcpALS classifier. Bars are color-coded according to the ranking in contribution for the given classifier, from dark-blue to white for high to low ranking. (**D**) Barplots showing SFPQ (black bars) and FUS (white bars) N/C ratios relative contribution to the different classifiers or PCs when applied. (**E**) Examples of motor neurons in the ventral spinal cord of wild-type, VCP^A232E^ and SOD1^G93A^ mice. Images were acquired as confocal z-stacks using a Zeiss 710 confocal microscope with a z-step of 1 μm, and processed to obtain a maximum intensity projection. MN cytoplasm is identified by ChAT immunofluorescence, nuclei are counterstained with DAPI, and SFPQ proteins are localised with immunofluorescence (green channel). For each MNs the disease probability is obtained either with comALS classifiers (green/red filled bars for LR/MLP classifiers respectively) or vcpALS classifiers (green/red empty bars for sLR and sMLP respectively). Scale bar = 26 μm.

We next analysed how each of the three categories of measurements, namely area-shape, texture and intensity, contributed to these classifiers using LMM (see Material and Method). We found that area-shape related measurements exhibited higher (although non significant) relative contribution to the *comALS* classifiers, while texture- and intensity-related measurements contributed more significantly to the *vcpALS* classifiers (**Fig. 4B**). Extraction of the five top contributors to either LR or sLR further revealed that the area-shape-related Zernike moments are indeed in the top five contributors of the *comALS* but not *vcpALS* classifiers (**Fig. 4C**), as further confirmed in MLP versus sMLP top five contributors (**Supplementary Fig. 4D**).

We previously showed that MNs from VCP-mutant mice exhibit reduced N/C ratios of FUS and SFPQ proteins and that SFPQ is also mislocalized in SOD1-mutant ALS models, while FUS is not (Luisier *et al*., 2018; Tyzack *et al*., 2019). Specifically looking at the relative contribution of SFPQ and FUS N/C ratios’ to the classifiers showed a stronger impact on the vcpALS classifiers compared to the comALS classifiers (**Fig. 4D** and **Supplementary Figs. 4E,F**). This corroborates our previous findings that showed increased protein mislocalization in VCP-mutant compared to SOD1-mutant ALS mouse models (Luisier *et al*., 2018; Tyzack *et al*., 2019). Visual inspection of randomly chosen tissue sections from example animals further confirmed that aberrant SFPQ protein mislocalization in VCP mutant MNs can easily be captured by eye and that SOD1- and VCP-mutant cells share features which are more challenging to pick up by inspection, hence reinforcing the importance of machine learning methods which can identify subtle changes from high-content data (**Fig. 4E**).

### Morphometric descriptors also capture phenotypes in human tissue sections from sporadic ALS cases

We next sought to validate our findings in human post-mortem tissue (PMT) from ALS patients. We previously analysed spinal cord sections from healthy and sporadic ALS (sALS) PMTs, immuno-labeled for either FUS or SFPQ, where nuclear and cytoplasmic compartments were manually identified with DAPI and ChAT (Luisier *et al*., 2018; Tyzack *et al*., 2019). Using these same images (**Tables S2-3**) and a combination of Ilastic and CellProfiler (see Materials and Methods and **Supplementary Figure 1B**), we automatically identified 432 MNs (66 stained for SFPQ and 366 stained for FUS) from which we extracted 850 morphological features (**Fig. 5A**). Unsupervised hierarchical clustering (euclidean distance and average clustering) using the morphological profiles across either the 66 MNs stained for SFPQ or the 366 MNs stained for FUS did not lead to clear segregation of the cells based on disease status of the human samples (**Fig. 5B**), suggesting that ALS phenotype is more subtle in human PMT compared to ALS mouse models. Nevertheless, SVD analysis of either the SFPQ data or FUS data, both rich-content data as shown by the high Shannon Entropies (**Supplementary Fig. 5A**), confirmed the presence of disease-related phenotype with the association of three principal components in either SFPQ or FUS data with sALS (**Supplementary Fig. 5B**).

**Figure 5.**
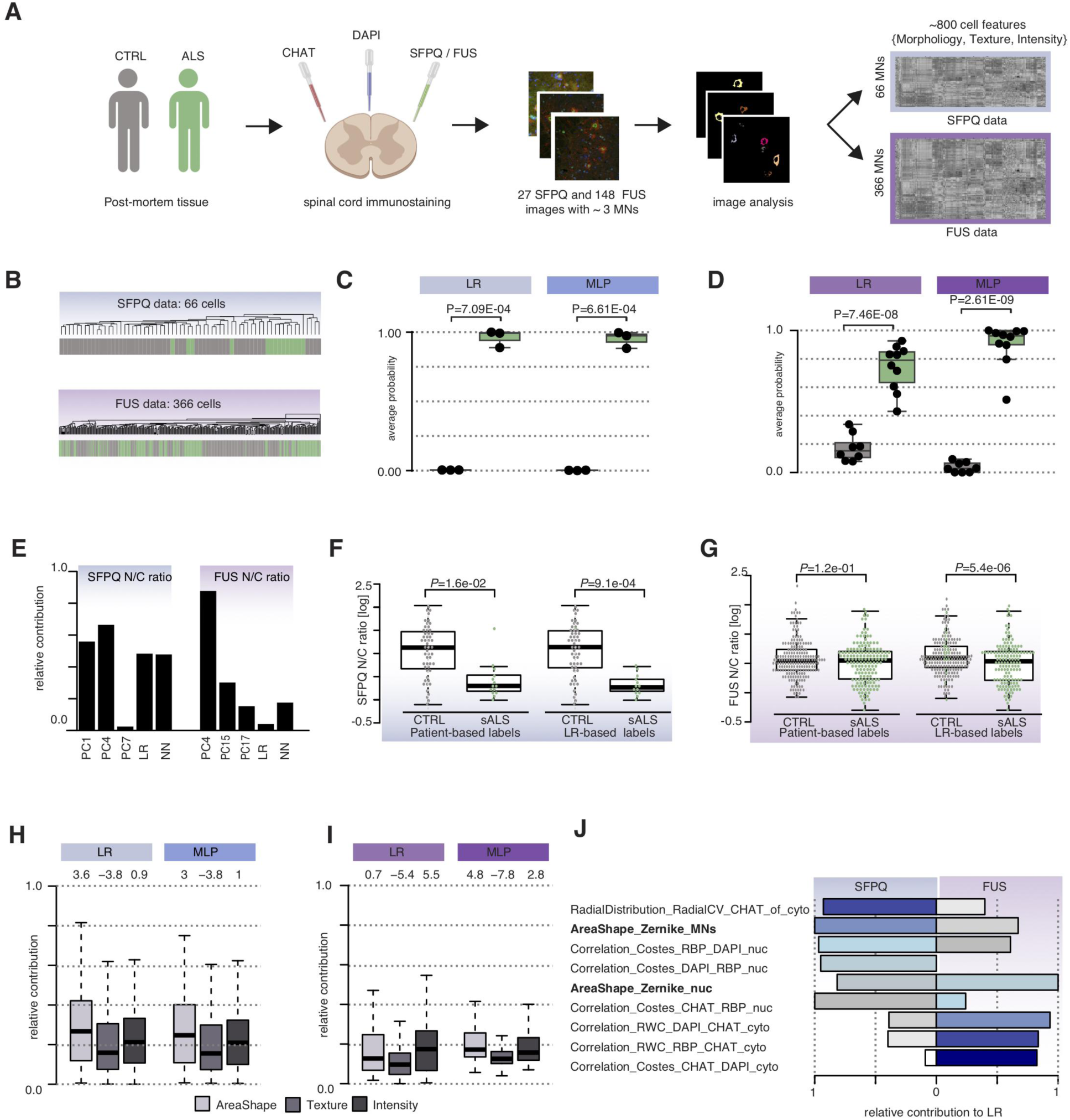
The Zernike moments capture aberrant cell behaviour in human ALS pathological post-mortem tissues. (**A**) Workflow for generating morphological profiles of MNs in ALS pathological spinal cord sections. Spinal cord sections from healthy donors and sporadic ALS patients were co-stained against ChAT, DAPI, SFPQ or FUS and then imaged (see **Table S2** for metadata details). A combination of *Ilastik* (Berg *et al*., 2019*a*) and *Cell Profiler* (Carpenter *et al*., 2006) enables automatic identification and feature extraction from cytoplasmic and nuclear compartments. (**B**) Unsupervised hierarchical clustering using 800 scaled measurements of the 66 cells co-immunolabeled with SFPQ or of the 366 cells co-immunolabeled with FUS do not show evidence for sALS phenotype. Grey bars = control MNs; green bars = sALS MNs. Euclidean distance and average clustering method. (**C**,**D**) The ability for LR and MLP clustering algorithms to detect the sALS effect in either SFPQ-stained data (**C**) or FUS-stained data (**D**) is assessed by comparing the disease probabilities of either the sALS group with those of the control group. The disease probability of each individual is obtained by using the mean probabilities of their cells to be sick according to 1) LR or 2) MLP classifiers. Data shown as box plots in which the centre line is the median, limits are the interquartile range and whiskers are the minimum and maximum. Dots are the individual disease profile. *P*-values obtained from Welch’s t test. (**E**) Barplots showing SFPQ (*left*) and FUS (*right*) N/C ratios relative contribution to the different classifiers or PCs when applied. (**F**) Boxplot showing the SFPQ nuclear/cytoplasmic (N/C) ratio per cell from three healthy donors and three sALS patients either grouped according to the patient-based labeling (*left*) or grouped according to the LR classifiers (*right*). Grey dots = control MNs; green dots = sALS-derived MNs. *P*-value obtained using Welch’s t test. (**G**) Boxplot showing the FUS nuclear/cytoplasmic (N/C) ratio per cell from eight healthy donors and ten sALS patients either grouped according to the patient-based labeling (*left*) or grouped according to the LR classifiers (*right*). Grey dots = control MNs; green dots = sALS-derived MNs. *P*-value obtained using Welch’s t test. (**H,I**) Boxplots showing the distributions of the relative contributions of area-shape (light grey), texture (grey) and intensity (dark grey) related measurements in LR and MLP classifiers either using SFPQ data (**H**) or FUS data (**I**). Z-scores obtained by a permutation test to assess the significance of the measurement category to each classifier are shown above each boxplot. Data shown as box plots in which the centre line is the median, limits are the interquartile range and whiskers are the minimum and maximum.(**J**) Barplots showing the relative contribution of the top five measurements in LR in either SFPQ or FUS data. Zernike moments either in the nucleus or the whole MNs contribute largely to both LR classifiers. Bars are color-coded according to the ranking in contribution for a given classifier, from dark blue to white for high to low ranking.

We next trained a combination of GMM with sALS related components, as well as LR and MLP for sALS automated identification. While the different methods showed very similar accuracy for the SFPQ data (**Supplementary Fig. 5C**), LR and MLP led to the highest accuracy for FUS data (**Supplementary Fig. 5D**). This would suggest that the sALS phenotype is supported by weighted linear combinations of all measurements in FUS data and that a restricted list of morphological profiles captured by two principal components is sufficient to capture cellular characteristics associated with sALS in SFPQ data. Further comparison of the abilities of each classifier to discriminate control from sALS groups using the per-individual probability to be sick confirmed our prior finding in mice that LR and MLP are the most efficient classifiers to capture sALS phenotype in high-content imaging data (**Figs. 5C,D** and **Supplementary Figs. 5E,F**). We next looked at the relative contribution of either SFPQ or FUS N/C ratio in these classifiers. We found that SFPQ localisation has comparatively more weight than FUS in discriminating sALS from control cells in LR and ML (**Fig. 5E**). This result is in line with our previous findings showing that, while both SFPQ and FUS proteins exhibit significant mislocalization in ALS, the extent of mislocalization is larger for SFPQ than for FUS(Luisier *et al*., 2018; Tyzack *et al*., 2019).

We next sought to assess the effect of MNs re-labelling on the significance of SFPQ and FUS protein mislocalization. We therefore performed Welsh’s t-test comparing the protein N/C ratios between two groups, either using the patient-based labels or the healthy versus disMNs labeling derived from the classifiers. Remarkably LR-based re-labeling decreased *P*-values in both data-sets, confirming the increase in signal-to-noise ratios upon re-classification of the cells (**Figs. 5F,G**).

Using a permutation test, we next tested the relationship between the relative contribution of each category of measurements (area shape, texture and intensity) in the classifiers that best discriminate sALS from control cells, namely LR and MLP. We found that area-shape related measurements significantly contribute to these models in both SFPQ and FUS data (**Figs. 5H,I**). Further looking at the five top relative contributors in LR in either data-sets revealed that area-shape-related Zernike moments are again among the top five measurements that contribute to the LR classifier in both SFPQ and FUS data (**Fig. 5J**), as well as in MLP classifiers (**Supplementary Fig. 5G**). Cumulatively, these findings suggest that common cellular disease phenotypes occur in human and mouse ALS models and are captured by the Zernike moments.

## DISCUSSION

### Novel pipeline for rapid processing and automated analysis of pathological sections

Our previous studies and those of others have relied on manual segmentation of pathological sections followed by analysis of single measurements. Here we developed a pipeline that couples fluorescence microscopy with highly quantifiable, automated and reproducible morphological profiling of MNs to allow fast and unbiased recovery of single-cell morphology profiles. By applying SVD analysis to the hundreds of cellular measurements derived from this platform we uncovered independent complex cellular phenotypes that associate with ALS MN populations. By showing that ALS phenotypes naturally emerge from such multichannel fluorescence microscopy high-dimensional data, we confirm the richness of the data over single-cell measurements and its relevance for future investigations.

Unbiased profiling at the single cell level represents a versatile and powerful readout for many cell states capturing the mechanistic details of a wide range of bioactivities (Feng *et al*., 2009; Ljosa *et al*., 2013; Wawer *et al*., 2014). ALS-related molecular events (e.g. mislocalization of an RBP from the nucleus to cytoplasm) may trigger events that lead to the divergence of outcomes amongst single cells. A recent study has indeed shown that there is substantial intrinsic heterogeneity at the single cell level among ALS cells (Tam *et al*., n.d.), which suggests that looking at the tissue as a whole might miss valuable insights into the underlying ALS pathobiology. Thus studies such as these highlight not only the importance of evaluating molecular phenotypes at the single cell level, but also the need to have appropriate tools and pipelines for the task. Here we show that machine learning methods such as LR and MLP capture combinations of subtle morphological alterations, without focusing on specific cell measurements, to accurately classify MNs in ALS and control groups. Remarkably the re-classification of MNs into healthy and sick cells using these label-independent classifiers led to an increase in signal-to-noise ratio for specific markers as well as an increase in significance in protein mislocalization. We also propose a combination of two single-cell disease scoring metrics derived from these classifiers that will be instrumental for future investigations of disease development using histopathological tissue sections.

Thus our study represents a significant advance in ALS histopathology as it presents an integrated pipeline for automatic identification and classification of MNs in pathological sections from raw image processing to the figures presented here. To facilitate the ready application and future development of methods for automatic identifications and scoring of ALS cells, we provide our complete raw image data sets, as well as open-source implementations of the various methods including the open-source workflow to carry out the segmentation, profiling and MNs automated classification.

### Morphometric descriptors capture common cellular phenotypes across human and mouse ALS models

Most studies related to ALS development report on molecular mechanisms that include aberrant protein homeostasis (ER stress and autophagy) and/or changes in RNA processing (Mackenzie *et al*., 2007; Zhang *et al*., 2015; Hall *et al*., 2017; Serio and Patani, 2018). Although cell morphology is intimately related to the physiological state of cells and intracellular mechanisms (Pincus and Theriot, 2007; Gomez *et al*., 2016; Ramkumar and Baum, 2016), there is not, to our knowledge, a comprehensive description of cell morphological changes in ALS. Here we show that area-shape related measurements, such as Zernike moments, can discriminate aberrant disease phenotypes across human and mouse ALS models. We also show that FUS and SFPQ N/C ratios have varying but lower weights in ALS MNs classifiers compared to the cellular characteristics described above. Zernike moments are the mapping of an image onto a set of complex orthogonal Zernike polynomials (Hwang and Kim, 2006) and have been proposed to provide a complete basis function representation of cell shape that preserves all the information about the shape of the cell (Mahi *et al*., 2014). These quantitative single cell shape descriptors have been shown to be reliable indicators of various cancers (Pincus and Theriot, 2007; Tahmasbi *et al*., 2011; Alizadeh *et al*., 2016). Our study suggests that area-shape measurements including Zernike moments may provide a means to read out the phenotypic state of ALS cells which cannot be easily captured by visual microscopic examination, hence the advantage of combining high-content microscopy data with machine learning methods to identify ALS sick cells. An important future avenue of this work will investigate the relationship between these area-shape measurements and intracellular molecular phenotypes.

### Future perspectives

Our work provides a framework for future studies with larger sample sizes, greater numbers of patients, and higher numbers of markers to rigorously test questions about the nature of heterogeneity between i) cells within a tissue and ii) patients. We envision that the ability to identify and interpret information contained in patterns of cellular heterogeneity in pathological sections will provide insights into physiology and disease that may be missed by traditional population-averaged or small-cell number studies. In particular, there are two specific areas where we envisage this method to be particularly informative for ALS, as well for other disorders. The first one is the study of the disease spatio-temporal evolution of the disease at the single cell level. In addition to progression in its severity, ALS also spreads spatially across the rostro-caudal axis (Wong *et al*., 2007). At present we do not have good data on what is the progression of changes that occur at the cell level, which could inform on the molecular pathways involved in spreading across anatomical regions. To address this we will be able to use the pipeline we developed here for a future study of serial sections along the spinal cord of individual animals to investigate spatio-temporal progression at different disease stages, as well as comparing the same anatomical regions of the spinal cord across the disease spectrum. This will offer a unique and crucial dataset on ALS progression. Another area where this method might be used, is in associating molecular phenotypes with cellular phenotypes at the single cell level. Specifically, by being able to reliably and automatically discriminate between cells that present a disease phenotype and cells that are still healthy (even within the same patient) we will be able to effectively create a framework for “digital molecular pathology” for ALS. Utilisation of these automated image analysis techniques can inform on which cells are i) most likely to be affected at any given time in individual patients, and therefore, ii) help to determine both the precise molecular pathomechanisms and potentially therapeutically treatable targets.

---

**Supplementary Figure 1.**
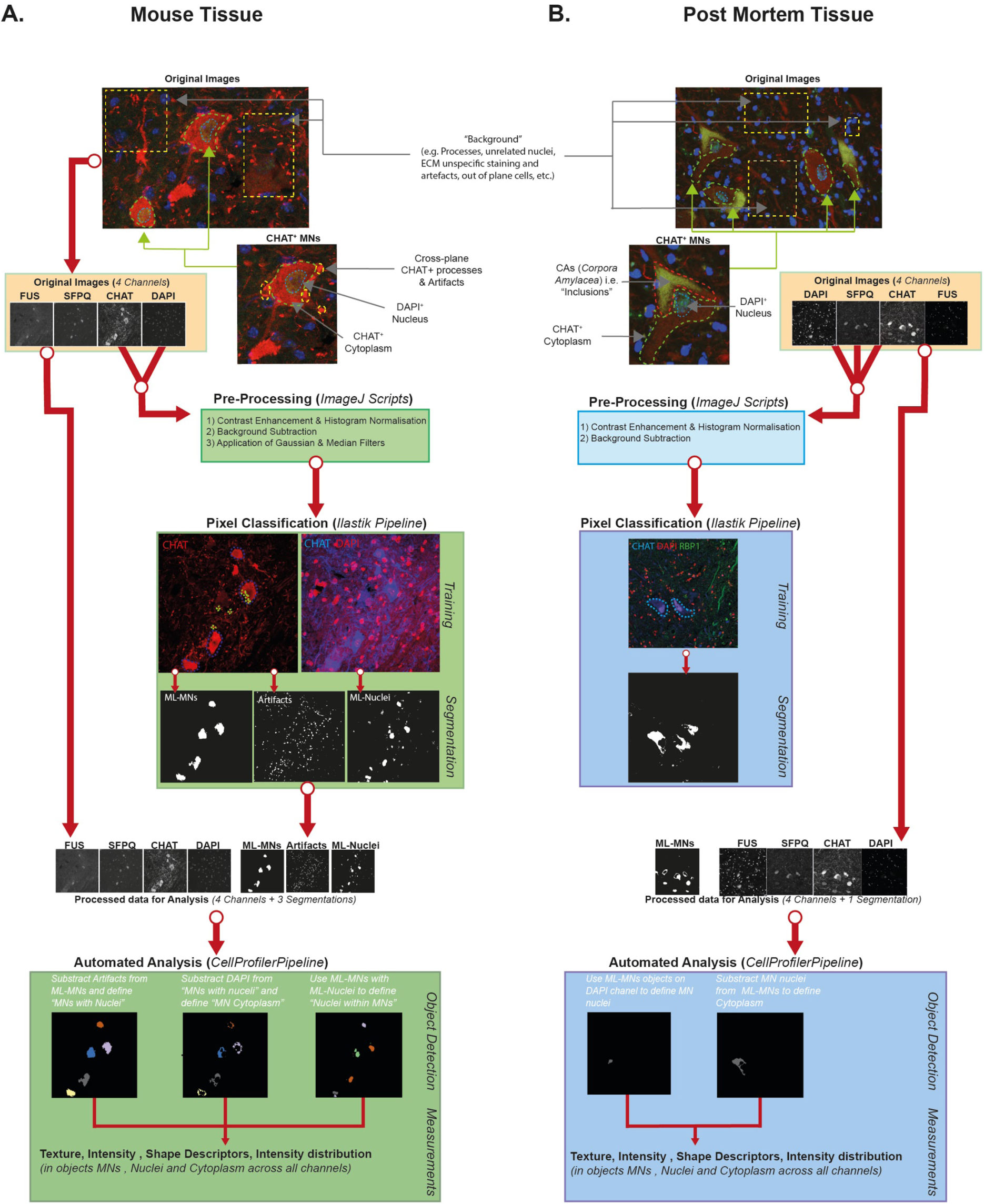
(**A**) Diagram representing the image processing workflow for mouse tissue immunolabeled for FUS, SFPQ, ChAT and counterstained with DAPI. To facilitate the MN segmentation, we first applied contrast enhancement, background correction and application of gaussian blur and median filters to ChAT stained images using ImageJ (Schneider *et al*., 2012). Next we used a subset of these preprocessed ChAT images to train a pixel classification algorithm in Ilastik (Berg *et al*., 2019*a*) for automated identification of artifacts and MNs. In parallel, automated nuclear segmentation was trained in Ilastik on randomly selected subsets of overlaid ChAT and DAPI images. Finally, all generated segmentations were added to the original dataset as additional channels, and used to identify the cytoplasm of MNs and perform automated densitometry and morphometric measurements in each compartment (cytoplasm, nucleus, whole MNs) in CellProfiler (Carpenter *et al*., 2006).(**B**) Diagram representing the image processing workflow for human post-mortem tissue immunolabeled for ChAT, FUS or SFPQ, and counterstained with DAPI. Individual channels were preprocessed in ImageJ for contrast enhancement and histogram equalisation. Next automated MN segmentation was trained using a subset of the original images using IIastik based on the three channels (SFPQ, DAPI, ChAT). Next DAPI channel and the masked MNs were provided to CellProfiler for automated segmentation of nuclei and cytoplasm followed by automatic acquisition of single-cell measurements in each compartment.

**Supplementary Figure 2.**
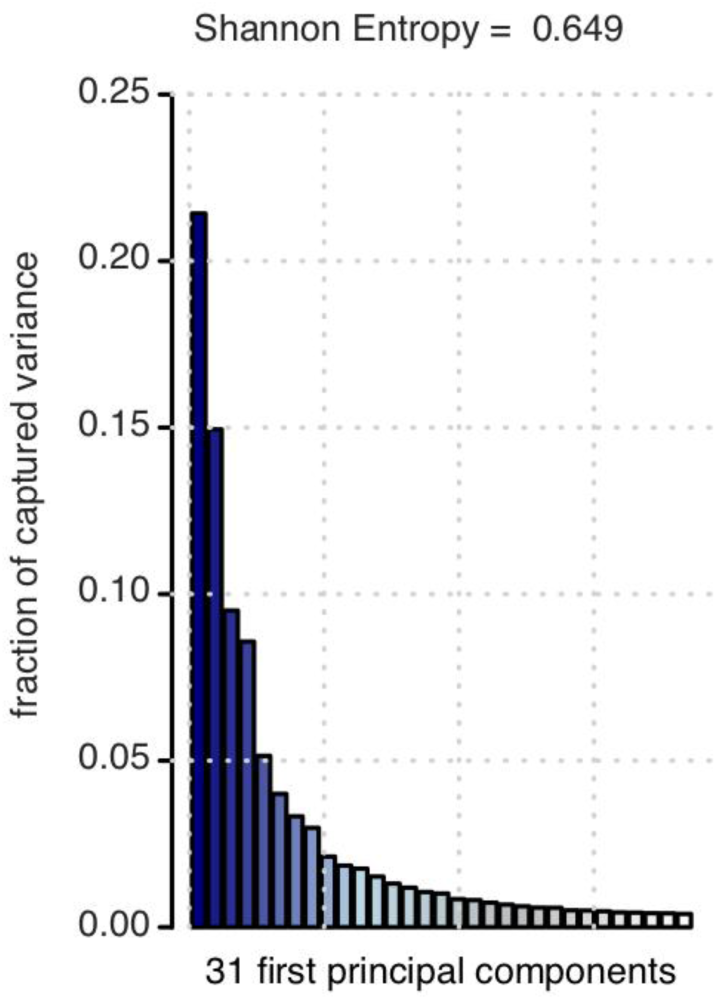
Fraction of explained variance captured by the first 31 principal components that captures 90% of the signal. Shannon Entropy of 0.65 indicates that the information in the data is well distributed among the principal components.

**Supplementary Figure 3.**
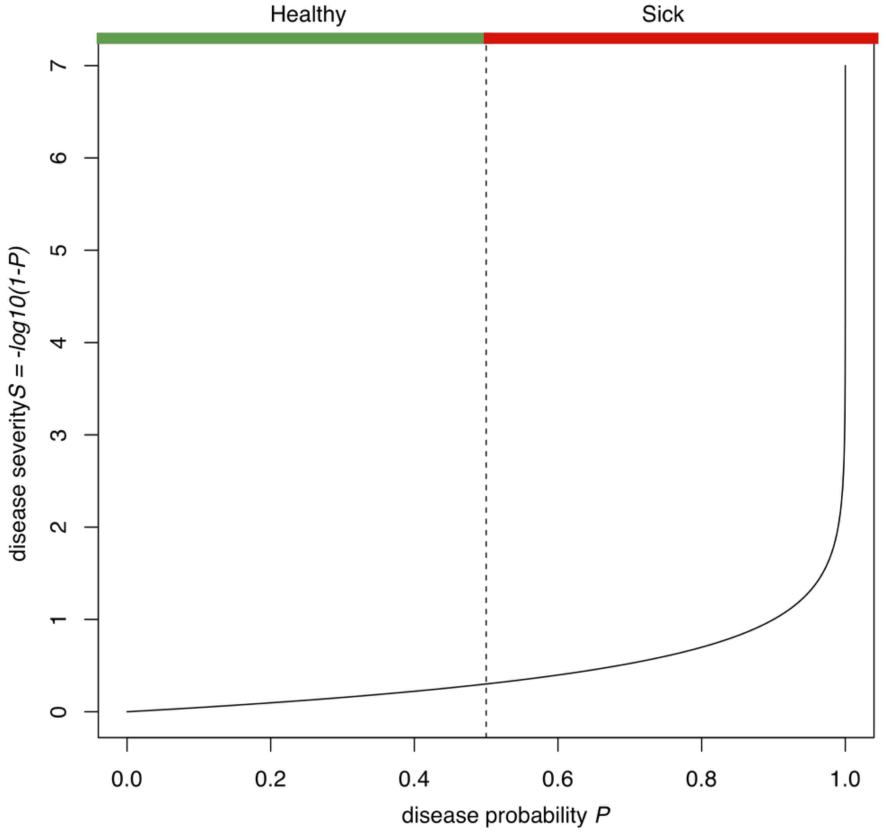
Comparison between disease probability P and disease severity S scores showing how similarly high disease probability can exhibit large differences in disease severity.

**Supplementary Figure 4.**
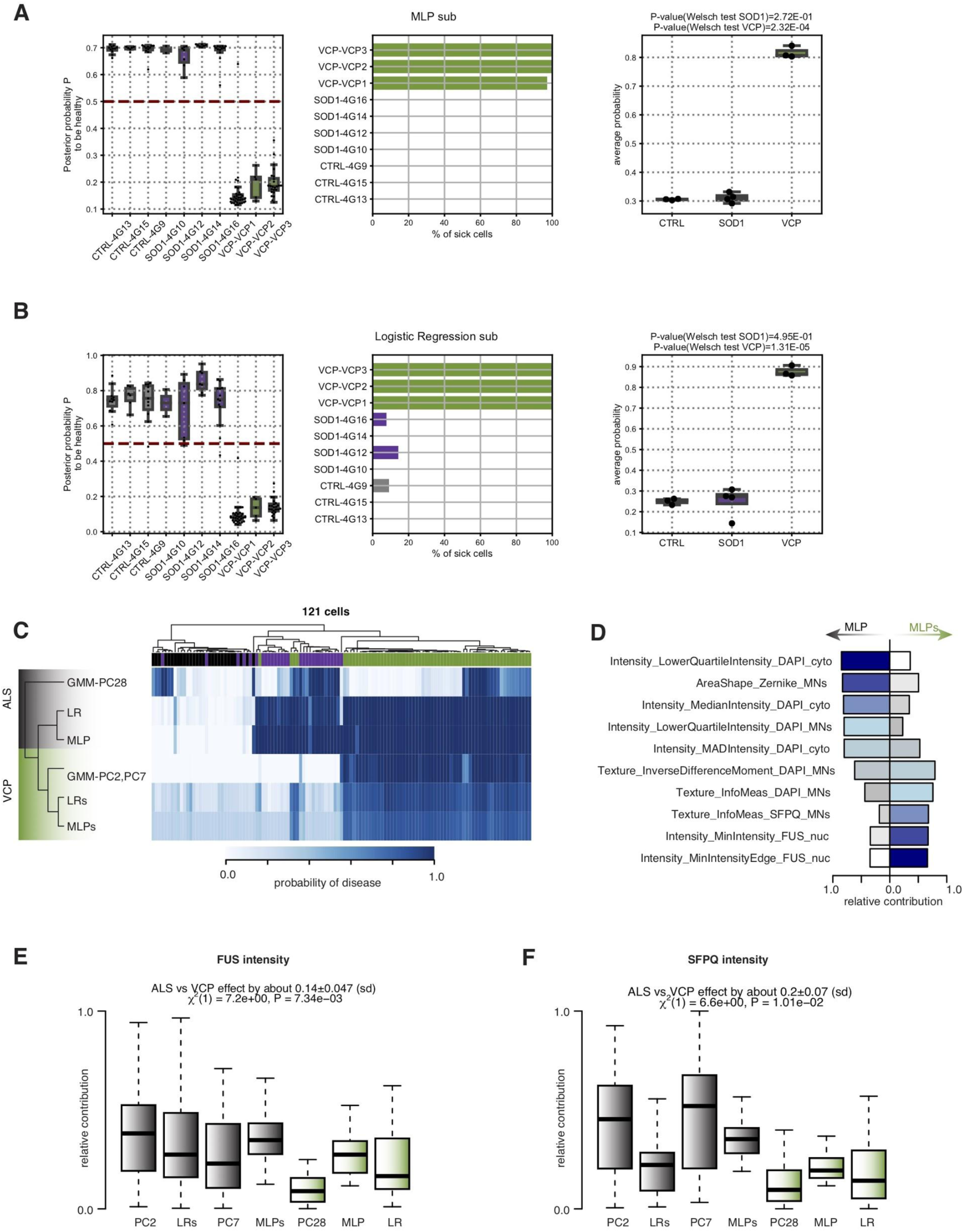
(**A**, **B**) MNs predicted probability distribution (*left*), per-animal percentage of sick cells (*center*), and per-animal disease probability *(right)* as obtained by LR classifier (**A**) and MLP classifier (**B**) trained on data censored for SOD1-mutant cells. (**C**) Heatmap showing the predicted disease probability for the 121 cells. Classifiers are hierarchically clustered using average on euclidean distances between disease probability profiles across the 121 cells. Green = *vcpALS* classifiers. Grey = *comALS* classifiers. (**D**) Barplots showing the relative contribution of the top five measurements in MLP and MLPs i.e. *comALS* versus *vcpALS* classifiers. Zernike moments either in the nucleus or the whole MNs contribute largely to ALS but not VCP classifier. Bars are color-coded according to the ranking in contribution for the given classifier, from dark blue to white for high to low ranking. (**E,F**) Boxplots showing the relative contribution of FUS (**E**) and SFPQ (**F**) intensity related measurements in *vcpALS* versus *comALS* classifiers. Linear mixed effects analysis of the relationship between the type of classifiers (*comALS* versus *vcpALS*) and the relative contribution of the measurement categories to account for idiosyncratic variation due to classifiers. Data shown as box plots in which the centre line is the median, limits are the interquartile range and whiskers are the minimum and maximum. Green = *vcpALS* classifiers; grey = *comALS* classifiers.

**Supplementary Figure 5.**
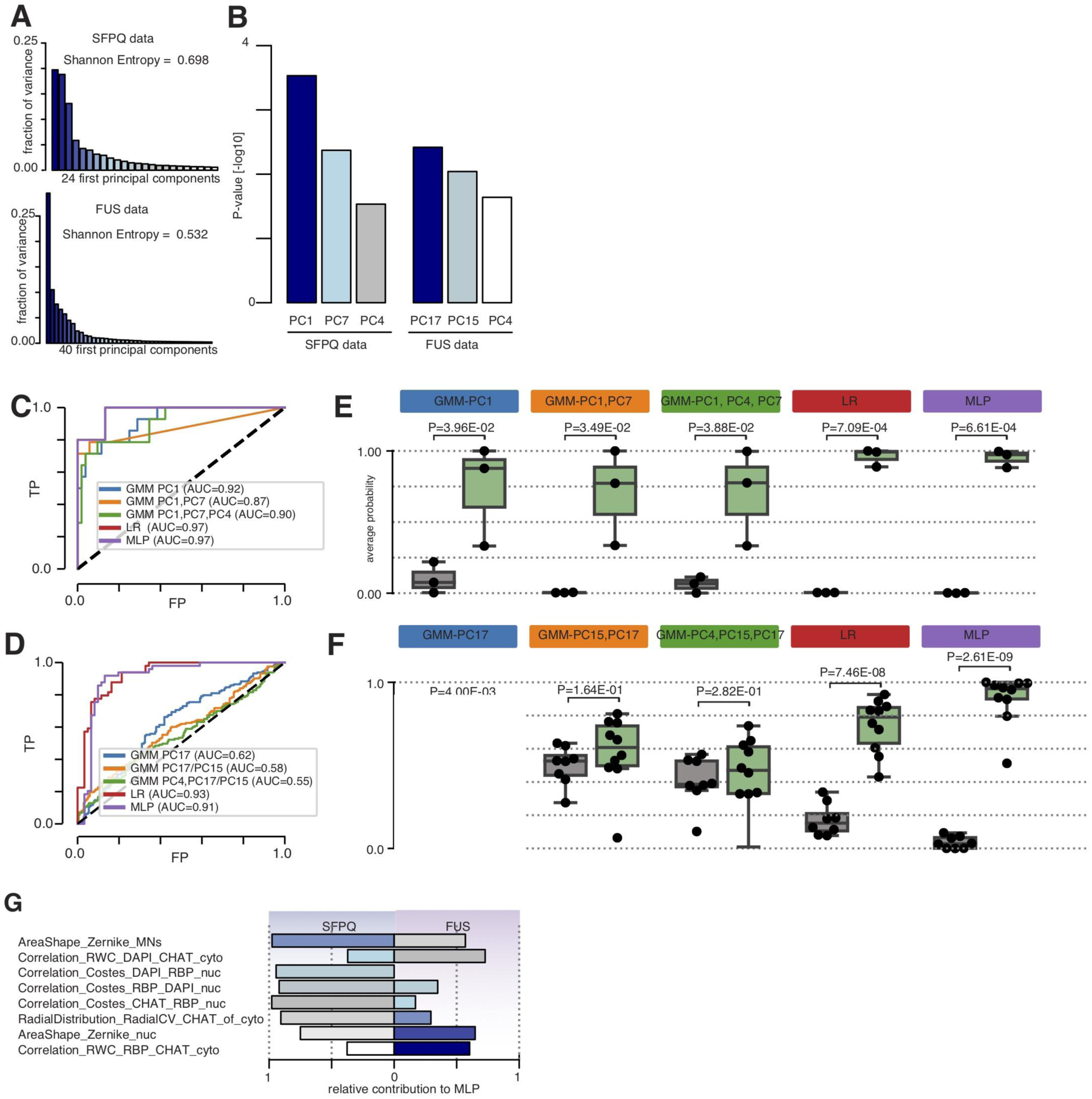
(**A**) Fraction of explained variance captured by the first 24 and 40 principal components that captures 90% of the signal in SFPQ and FUS data respectively. (**B**) Barplots showing the association between principal components and ALS in SFPQ (*left)* and FUS (*right*) data. Linear mixed effects analysis of the relationship between ALS phenotype and each of the 24 and 40 first principal components to account for idiosyncratic variation due to individuals shows significant association of PC1, PC4 and PC7 and ALS in SFPQ data, and association between PC4, PC15 and PC17 with ALS in FUS data. (**C**,**D**) Performance analysis of each classifier in SFPQ data (**C**) or FUS data (**D**) in their ability to discriminate sALS MNs from healthy MNs using receiver operating characteristic (ROC) curves and aurea under the curves (AUC). (**E**, **F**) The ability for each clustering algorithm to detect sALS effect is assessed by comparing the disease probabilities of sALS group with those of the control group. The disease probability of each individual is obtained by using the mean probabilities of its cells to be sick according to individual classifiers in SFPQ data (**B**) or FUS data (**F**). Data shown as box plots in which the center line is the median, limits are the interquartile range and whiskers are the minimum and maximum. Dots are the individual disease profile. *P*-values obtained from Welch’s t test. (**G**) Barplots showing the relative contribution of the top five measurements in MLP in either SFPQ or FUS post-mortem tissue data. Zernike moments either in the nucleus or the whole MNs contribute largely to both MLP classifiers. Bars are color-coded according to the ranking in contribution for the given classifier, from dark blue to white for high to low ranking.

**Table S1.** List of images used for FUS and SFPQ cellular localisation in (Luisier *et al*., 2018; Tyzack *et al*., 2019); mouse data.

| imageID | mutation | RBP1 | RBP2 | animal | origin | Lab | pairs |
| --- | --- | --- | --- | --- | --- | --- | --- |
| 4G10D1 | SOD1 | SFPQ | FUS | 4G10 | Lab_2 | Lab_2 | SFPQ_FUS |
| 4G10D2 | SOD1 | SFPQ | FUS | 4G10 | Lab_2 | Lab_2 | SFPQ_FUS |
| 4G12D1 | SOD1 | SFPQ | FUS | 4G12 | Lab_2 | Lab_2 | SFPQ_FUS |
| 4G12D2 | SOD1 | SFPQ | FUS | 4G12 | Lab_2 | Lab_2 | SFPQ_FUS |
| 4G13D1 | CTRL | SFPQ | FUS | 4G13 | Lab_2 | Lab_2 | SFPQ_FUS |
| 4G13D2 | CTRL | SFPQ | FUS | 4G13 | Lab_2 | Lab_2 | SFPQ_FUS |
| 4G13D3 | CTRL | SFPQ | FUS | 4G13 | Lab_2 | Lab_2 | SFPQ_FUS |
| 4G14D1 | SOD1 | SFPQ | FUS | 4G14 | Lab_2 | Lab_2 | SFPQ_FUS |
| 4G14D2 | SOD1 | SFPQ | FUS | 4G14 | Lab_2 | Lab_2 | SFPQ_FUS |
| 4G14D3 | SOD1 | SFPQ | FUS | 4G14 | Lab_2 | Lab_2 | SFPQ_FUS |
| 4G15D1 | CTRL | SFPQ | FUS | 4G15 | Lab_2 | Lab_2 | SFPQ_FUS |
| 4G15D2 | CTRL | SFPQ | FUS | 4G15 | Lab_2 | Lab_2 | SFPQ_FUS |
| 4G16D1 | SOD1 | SFPQ | FUS | 4G16 | Lab_2 | Lab_2 | SFPQ_FUS |
| 4G16D2 | SOD1 | SFPQ | FUS | 4G16 | Lab_2 | Lab_2 | SFPQ_FUS |
| 4G16D3 | SOD1 | SFPQ | FUS | 4G16 | Lab_2 | Lab_2 | SFPQ_FUS |
| 4G9D1 | CTRL | SFPQ | FUS | 4G9 | Lab_2 | Lab_2 | SFPQ_FUS |
| 4G9D2 | CTRL | SFPQ | FUS | 4G9 | Lab_2 | Lab_2 | SFPQ_FUS |
| 4G9D3 | CTRL | SFPQ | FUS | 4G9 | Lab_2 | Lab_2 | SFPQ_FUS |
| VCP1AFITC | VCP | SFPQ | FUS | VCP1 | Lab_1 | Lab_1 | SFPQ_FUS |
| VCP1BCY5 | VCP | SFPQ | FUS | VCP1 | Lab_1 | Lab_1 | SFPQ_FUS |
| VCP1BFITC | VCP | SFPQ | FUS | VCP1 | Lab_1 | Lab_1 | SFPQ_FUS |
| VCP1CFITC | VCP | SFPQ | FUS | VCP1 | Lab_1 | Lab_1 | SFPQ_FUS |
| VCP1DFITC | VCP | SFPQ | FUS | VCP1 | Lab_1 | Lab_1 | SFPQ_FUS |
| VCP1EFITC | VCP | SFPQ | FUS | VCP1 | Lab_1 | Lab_1 | SFPQ_FUS |
| VCP2A | VCP | SFPQ | FUS | VCP2 | Lab_1 | Lab_1 | SFPQ_FUS |
| VCP2AMODIFIED | VCP | SFPQ | FUS | VCP2 | Lab_1 | Lab_1 | SFPQ_FUS |
| **VCP2BFITC** | VCP | SFPQ | FUS | VCP2 | Lab_1 | Lab_1 | SFPQ_FUS |
| **VCP2BMODIF** | VCP | SFPQ | FUS | VCP2 | Lab_1 | Lab_1 | SFPQ_FUS |
| **VCP2CFITC** | VCP | SFPQ | FUS | VCP2 | Lab_1 | Lab_1 | SFPQ_FUS |
| **VCP2DFITC** | VCP | SFPQ | FUS | VCP2 | Lab_1 | Lab_1 | SFPQ_FUS |
| **VCP2EFITC** | VCP | SFPQ | FUS | VCP2 | Lab_1 | Lab_1 | SFPQ_FUS |
| **VCP2FFITC** | VCP | SFPQ | FUS | VCP2 | Lab_1 | Lab_1 | SFPQ_FUS |
| **VCP3A** | VCP | SFPQ | FUS | VCP3 | Lab_1 | Lab_1 | SFPQ_FUS |
| **VCP3B** | VCP | SFPQ | FUS | VCP3 | Lab_1 | Lab_1 | SFPQ_FUS |
| **VCP3C** | VCP | SFPQ | FUS | VCP3 | Lab_1 | Lab_1 | SFPQ_FUS |
| **VCP3D** | VCP | SFPQ | FUS | VCP3 | Lab_1 | Lab_1 | SFPQ_FUS |
| **VCP3E** | VCP | SFPQ | FUS | VCP3 | Lab_1 | Lab_1 | SFPQ_FUS |
| **VCP3F** | VCP | SFPQ | FUS | VCP3 | Lab_1 | Lab_1 | SFPQ_FUS |

**Table S2.**
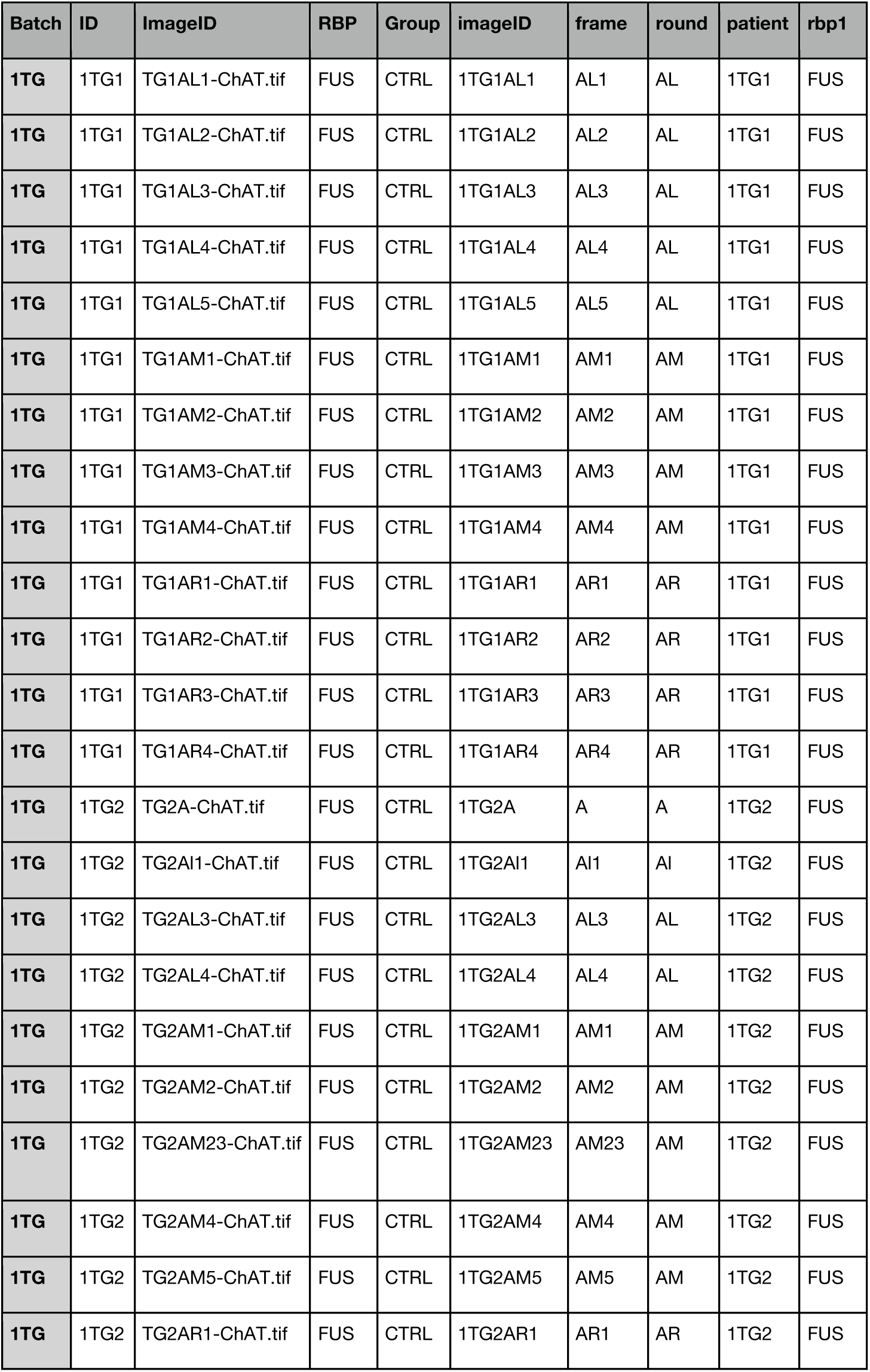
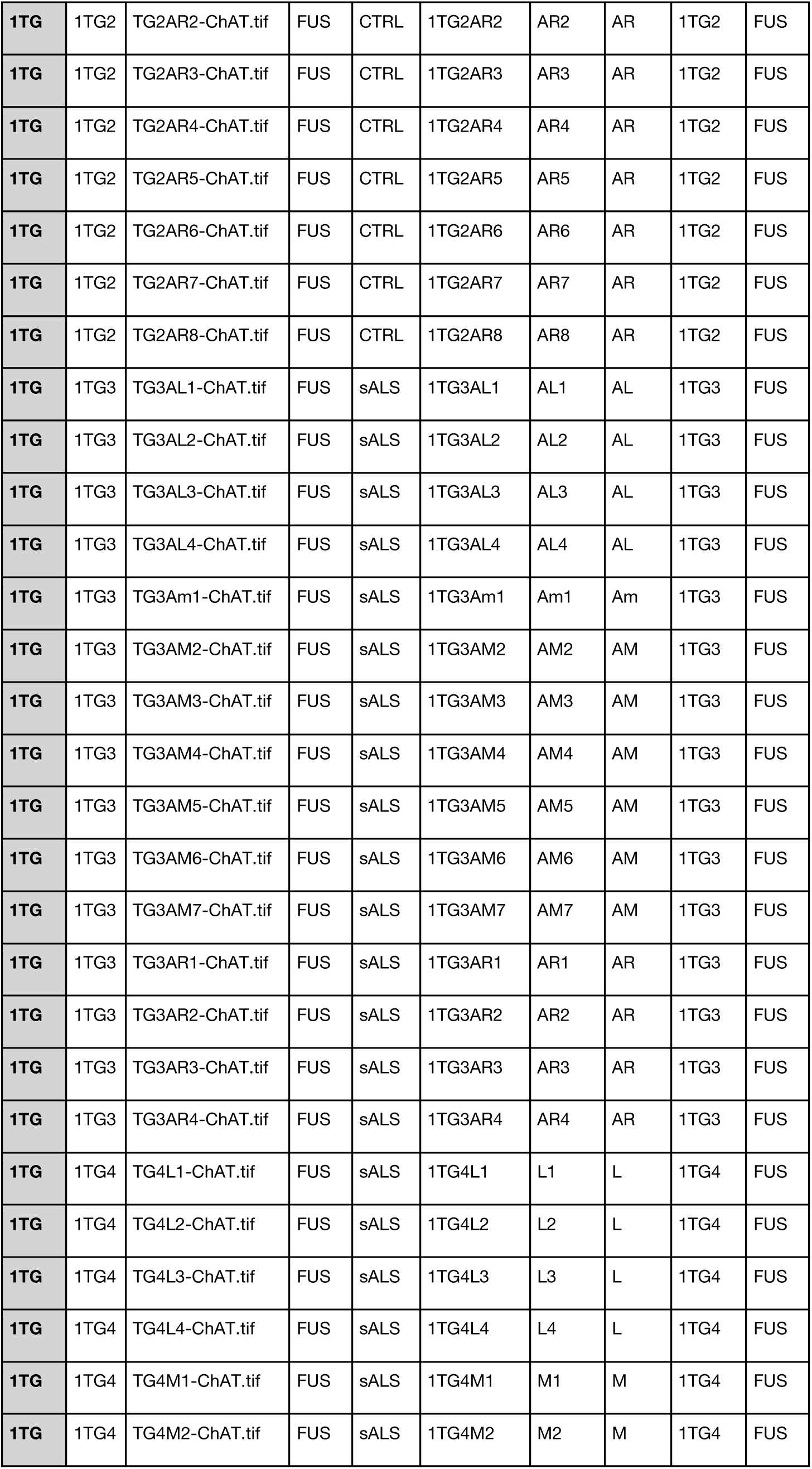
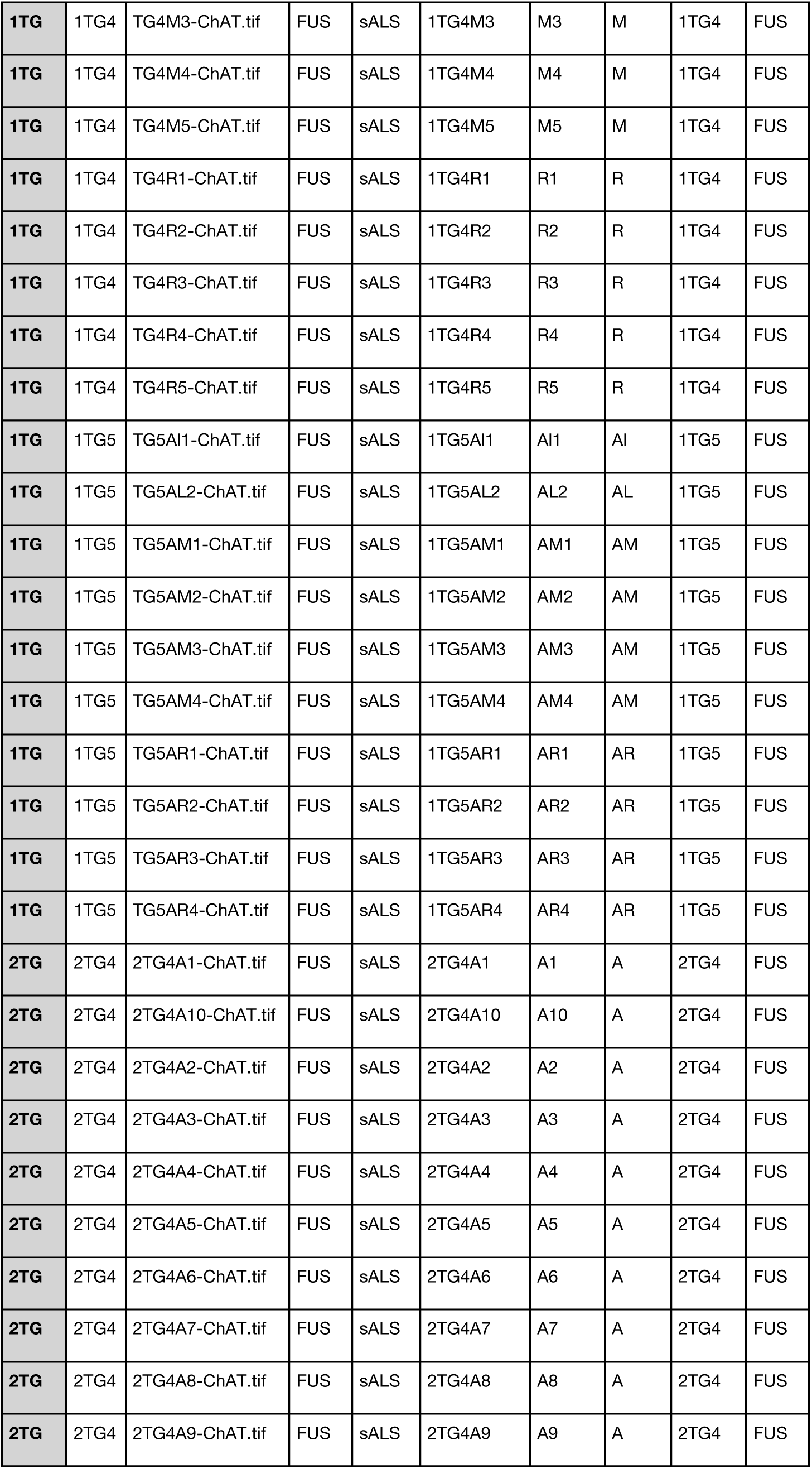
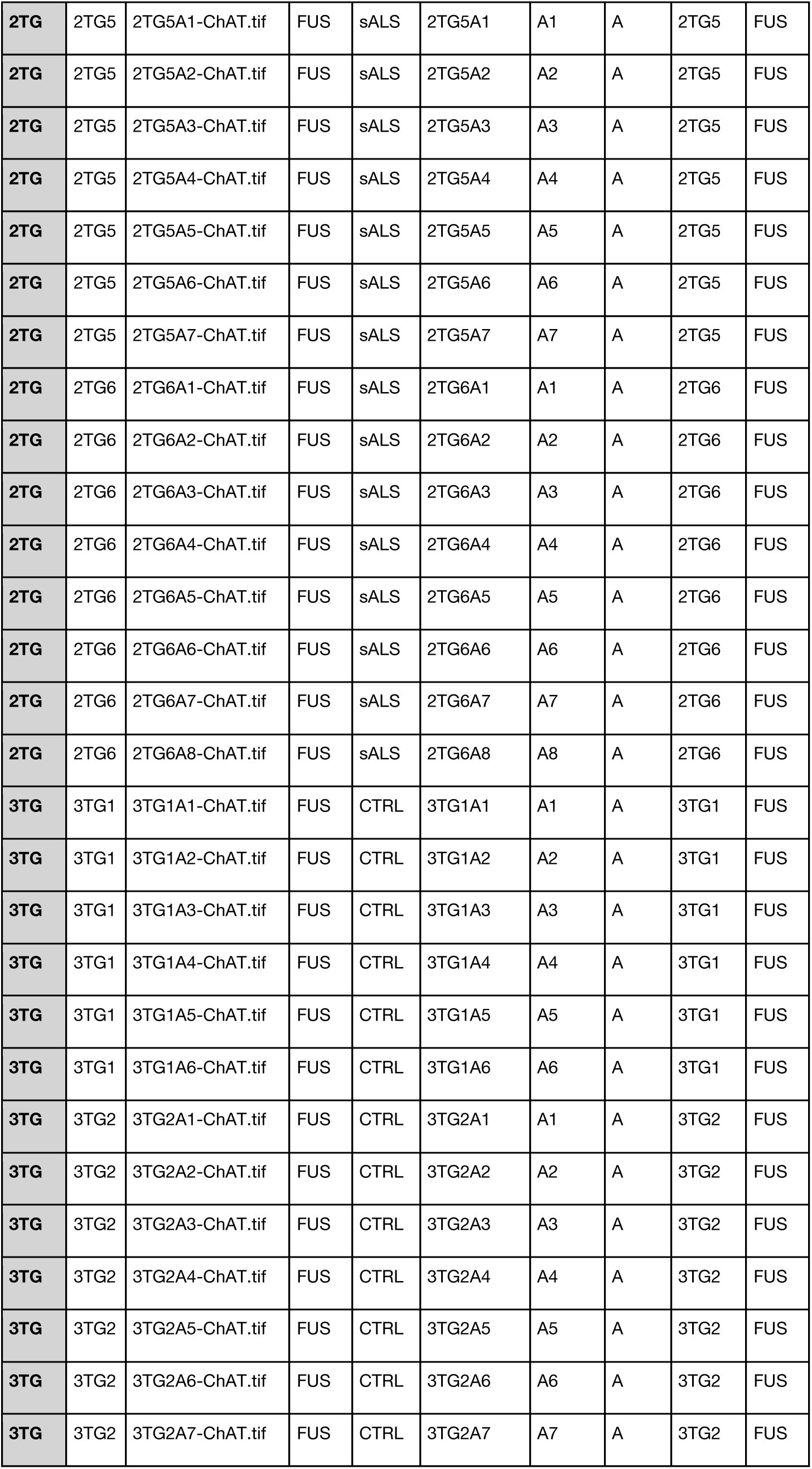
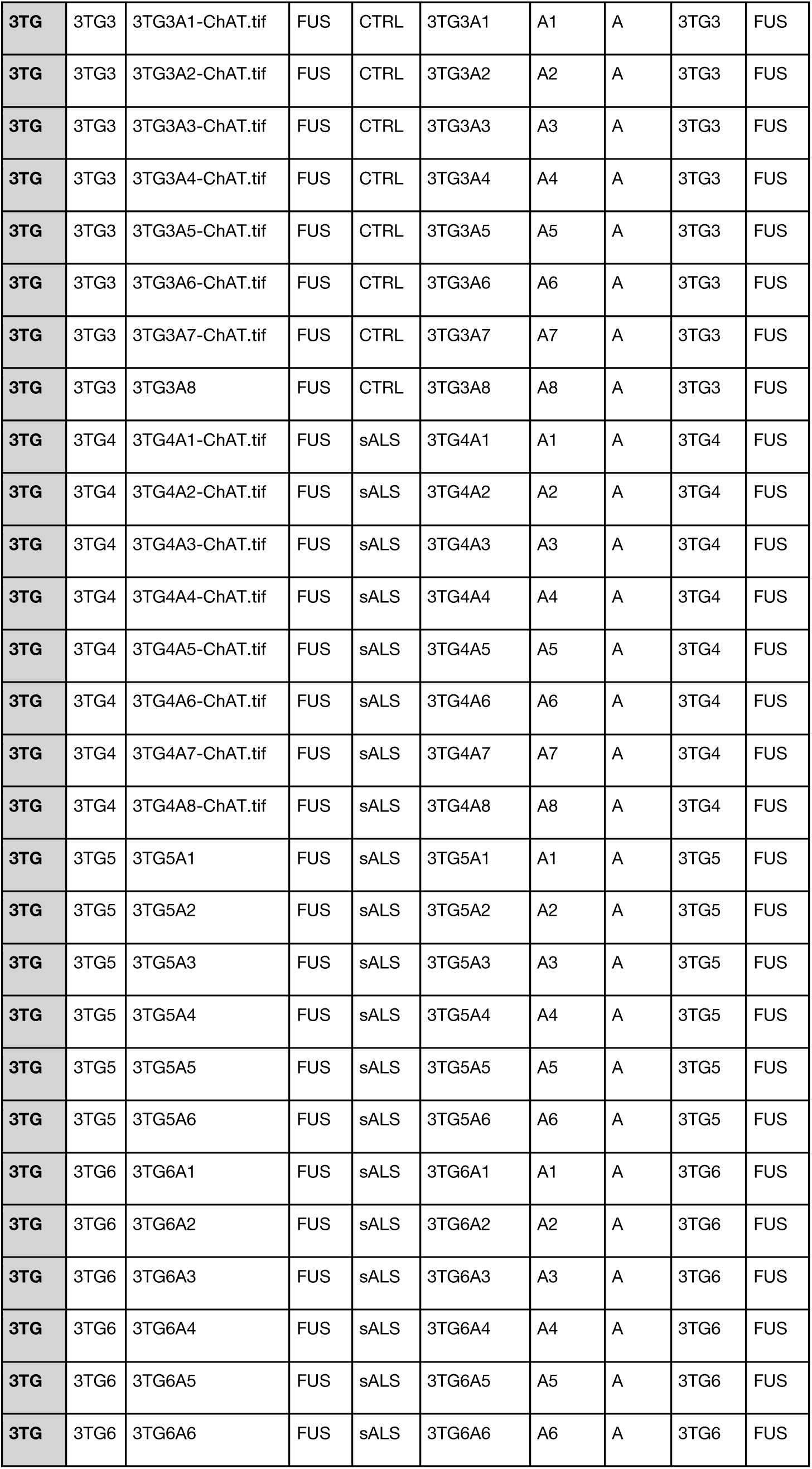
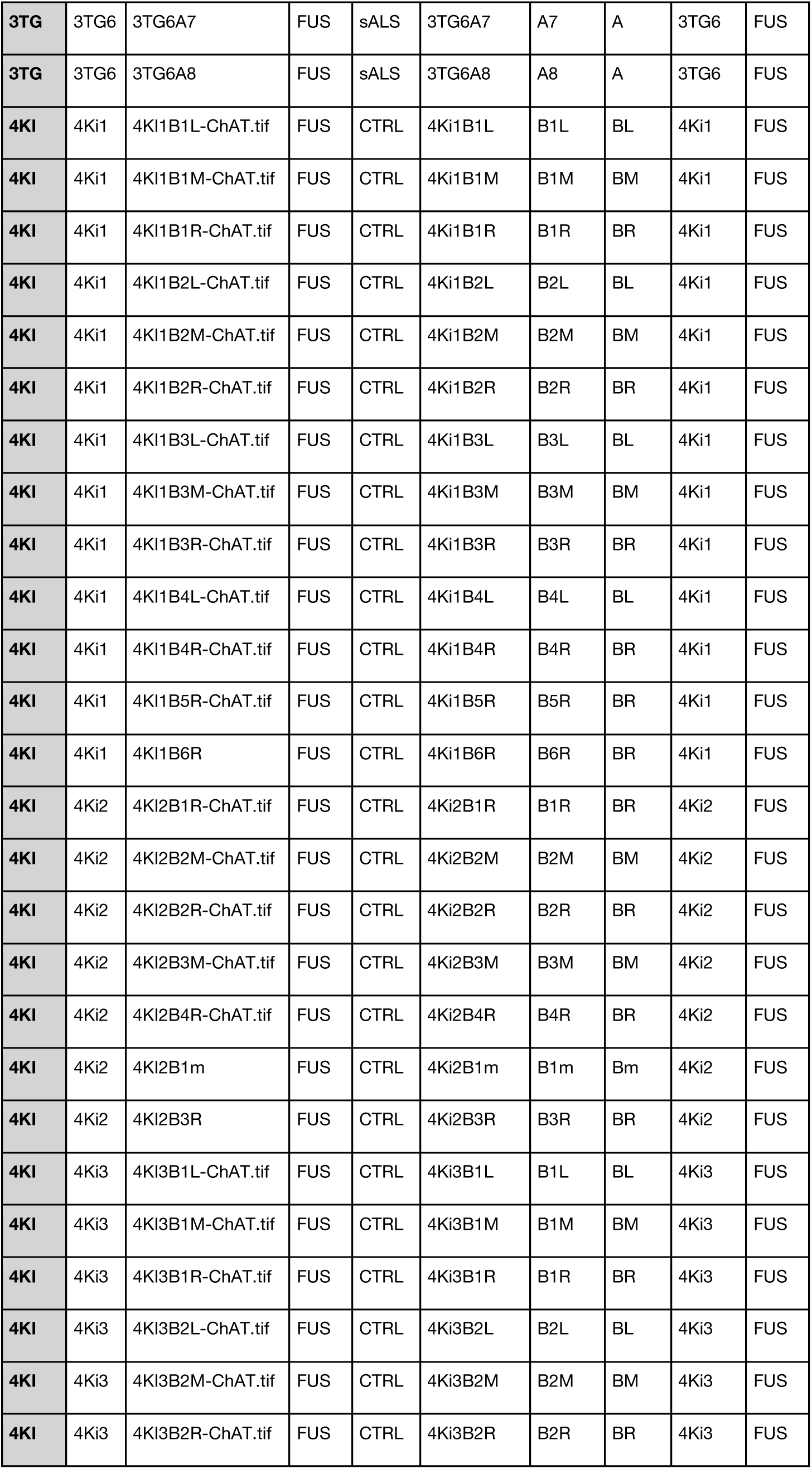
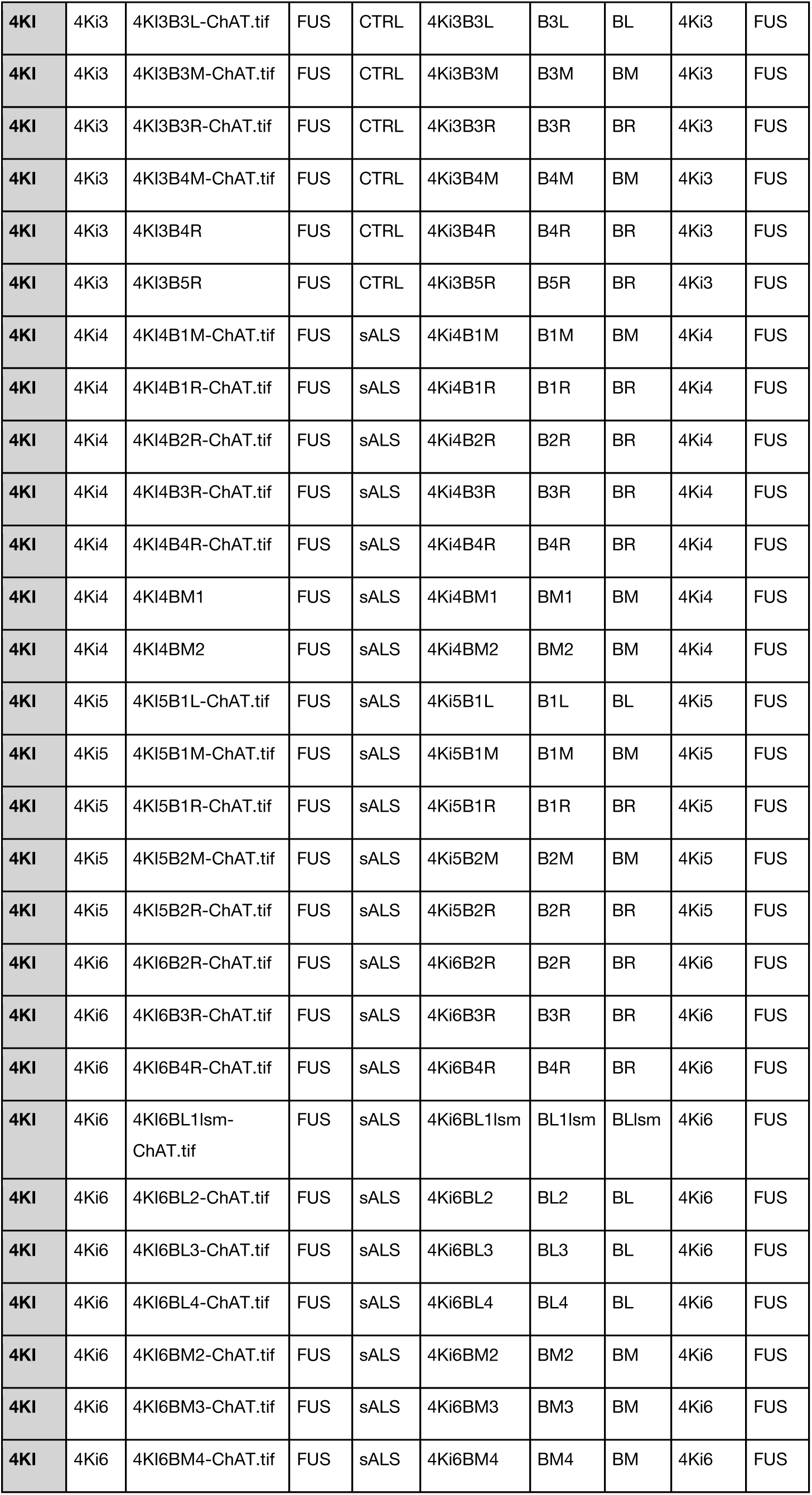
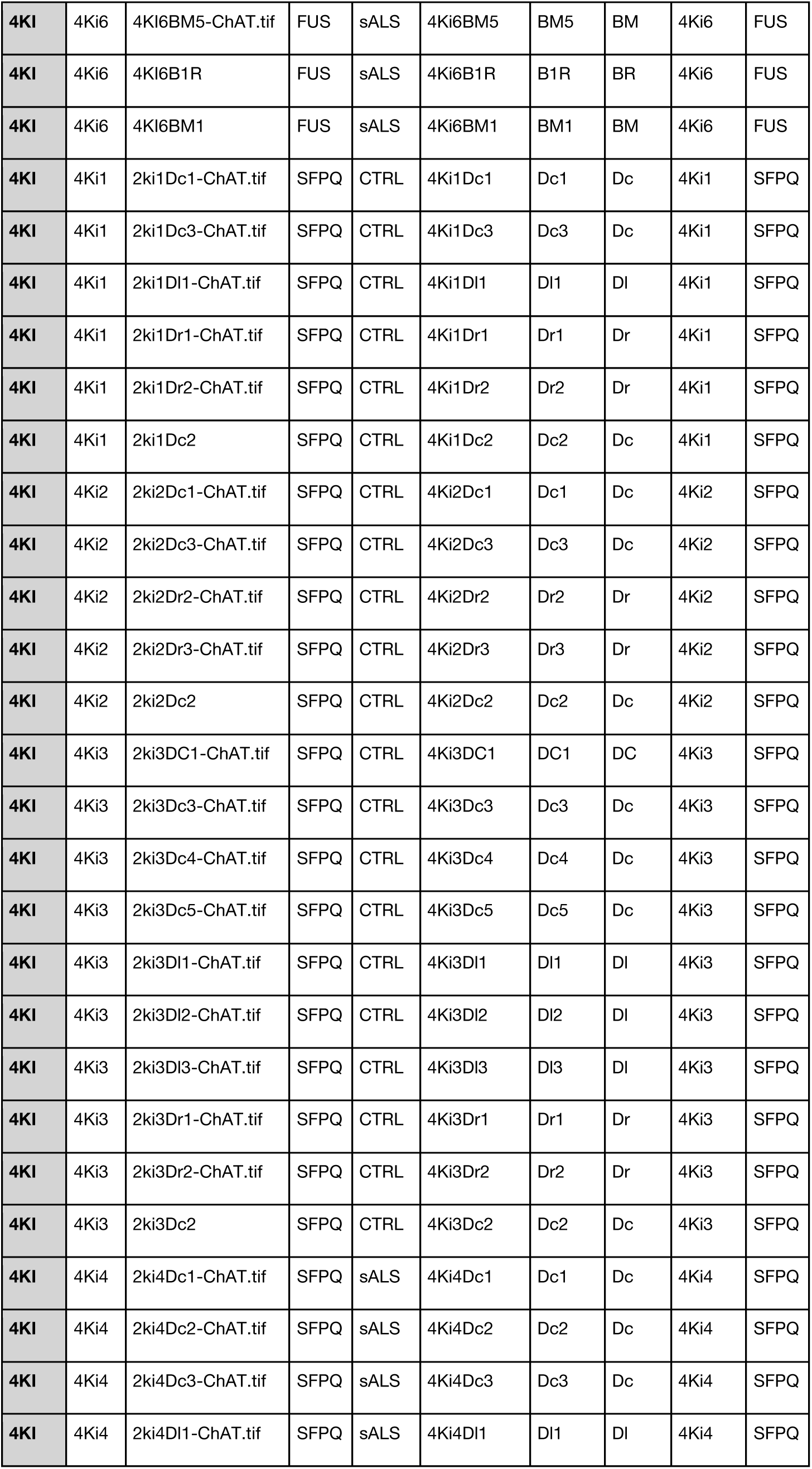
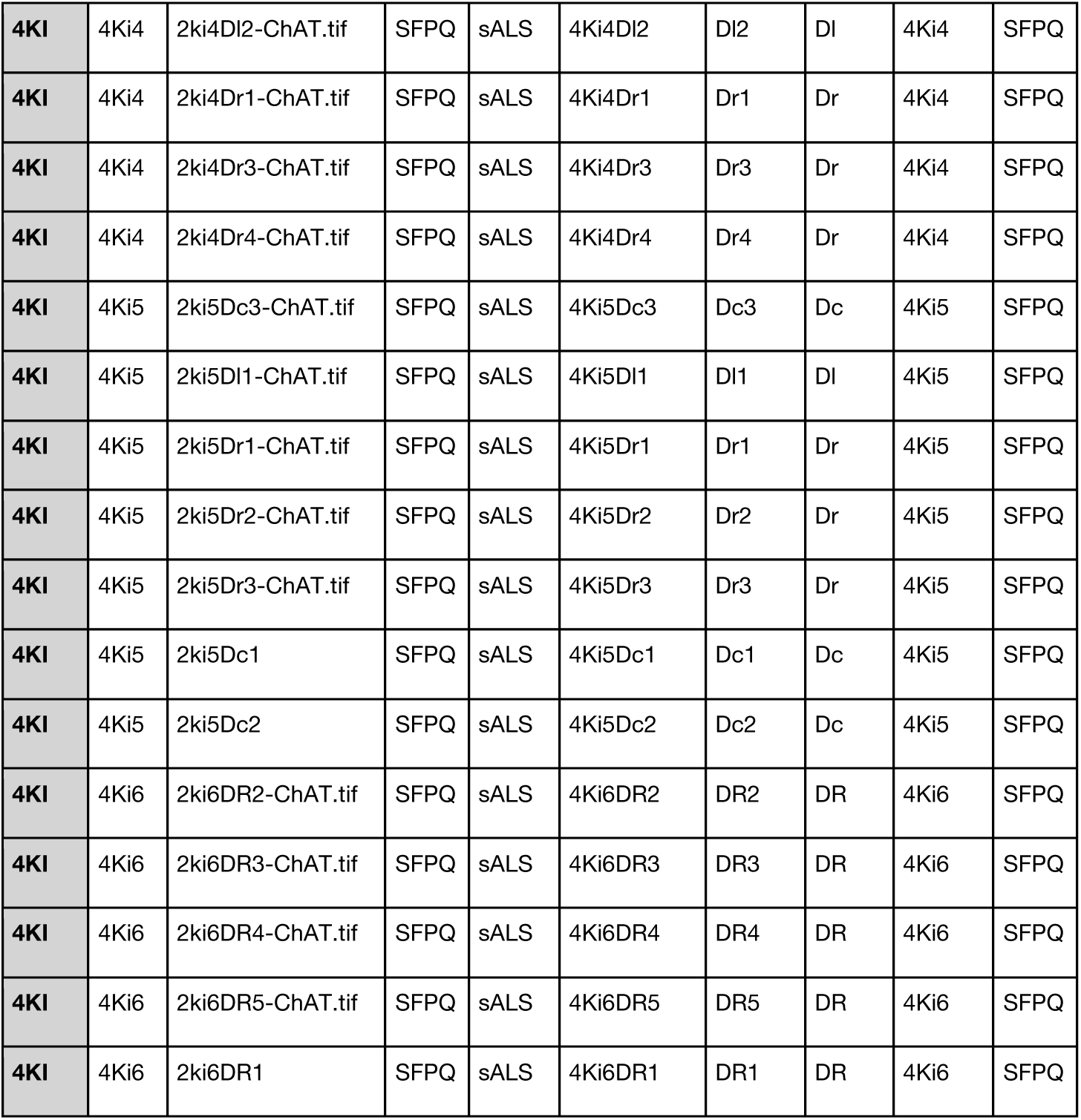
List of images used for FUS and SFPQ cellular localisation in (Luisier *et al*., 2018; Tyzack *et al*., 2019); human data.

**Table S3.**
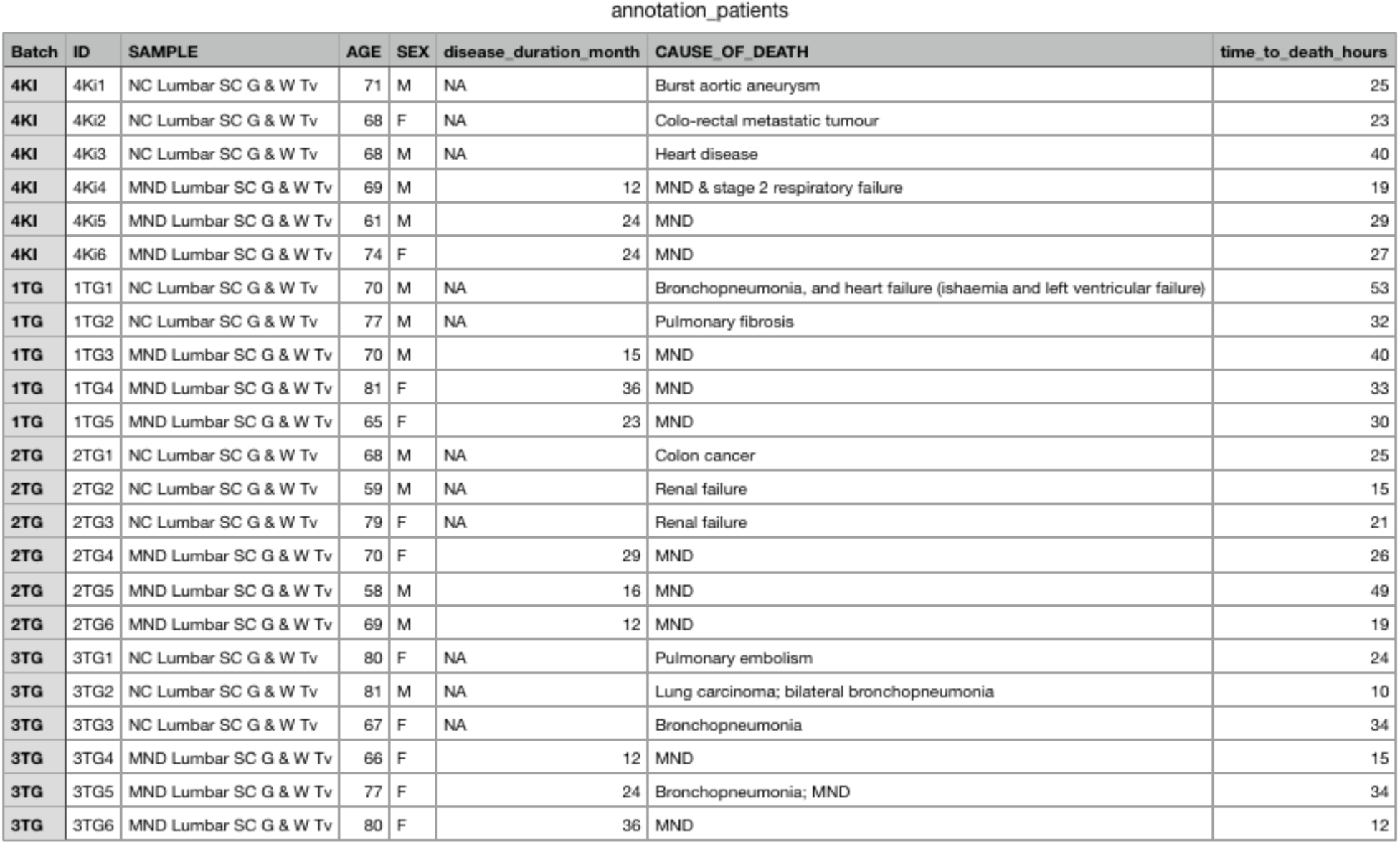
Description of the donors from which PMTs were obtained.

## MATERIALS AND METHODS

### Compliance with ethical standards

Experimental protocols were all carried out according to approved regulations and guidelines by the UCLH’s National Hospital for Neurology and Neurosurgery and UCL Queen Square Institute of Neurology joint research ethics committee (09/0272). The human post-mortem spinal cord samples were obtained from the tissue bank NeuroResource, UCL Queen Square Institute of Neurology, London, UK. Samples were donated to the tissue bank with written tissue donor informed consent following ethical review by the NHS NRES Committee London–Central and stored under a Research Sector Licence from the UK Human Tissue Authority (HTA). All animal experiments described in this study were carried out under Licence from the UK Home Office, and were approved by the Ethical Review Panel of the Institute of Neurology.

### Animals, transgenic models and tissue processing

All experiments were carried out following the guidelines of the UCL Institute of Neurology Genetic Manipulation and Ethic Committees and in accordance with the European Community Council Directive of November 24, 1986 (86/609/EEC). Animal experiments were undertaken under a Licence from the UK Home Office in accordance with the Animals (Scientific Procedures) Act 1986 (Amended Regulations 2012) and were approved by the Ethical Review Panel of the Institute of Neurology. The human spinal cord samples were obtained from the tissue bank NeuroResource, UCL Institute of Neurology, London, UK. Samples were donated to the tissue bank with written tissue donor informed consent following ethical review by the NHS NRES Committee London– Central and stored under a Research Sector Licence from the UK Human Tissue Authority (HTA). Animals, transgenic models and tissue processing were performed as in (Luisier *et al*., 2018; Tyzack *et al*., 2019). Indeed these data are utilised in the current manuscript and no additional animal usage was required.

### Automated image processing

Both mouse tissues and patient post-mortem samples were sectioned and immunostaining as described in our previous studies (Luisier *et al*., 2018; Tyzack *et al*., 2019). Raw images were processed by combining the following open source softwares (and packages) for automatic segmentation and measurements of cell features: ImageJ, (Schneider *et al*., 2012), Ilastik (Berg *et al*., 2019*a*)and Cellprofiler (Carpenter *et al*., 2006) Diagrams of the mouse and human pipelines are shown in **Supplementary Fig.1**.

#### Mouse Tissue pipeline (Supplementary Fig. 1A)

*Preprocessing:* choline acetyltransferase (ChAT) immunolabeling is a reliable MN marker used hereused for MN segmentation. To facilitate the segmentation, we first enhanced the contrast, subtracted the background and applied gaussian and median filters to each individual image, using the base scripts provided within the main FiJi (ImageJ) release(Schneider *et al*., 2012). Additionally we used the particle detection and image calculator packages in ImageJ to remove objects smaller than 2000px, as these were considered staining artifacts. A custom script combining these preprocessing steps in ImageJ is accessible in <u>Zenodo</u> under the accession number 3985099. Notably the modified images were only used for segmentation while original images were used for cell profiling.

*Pixel classification:* a randomly selected subset of the preprocessed images (ranging from 8 to 15) was used for training using a parallel random forest (VIGRA) algorithm in Ilastik (see labels and training set examples provided in Zenodo) for automated MNs segmentation.. For nuclei segmentation, a subset of DAPI and ChAT fluorescent images were next combined and training was undertaken in Ilastik (see labels and training set examples provided in Zenodo).

*Automated image analysis:* the three binary masks generated by the pixel classification algorithm (MNs, nuclei, inclusions) were added to the original channels and then used as a dataset for automated analysis in Cellprofiler (see pipeline provided in Zenodo). Here the masks were converted to objects for further processing. The inclusion-objects were subtracted from MN-objects and nuclei-objects to exclude measurements of staining artifacts. Then the resulting ‘clean’ nuclei-objects were used to detect MNs including nuclei. This is a crucial step for processing sliced images where nuclei can be physically separated from their corresponding MN. We continued processing these selected MNs by subtracting the nuclei-object and therefore defining the cytoplasm of this cell. As a final step, for each MN, texture, intensity, intensity distribution, size and shape were measured in MN-, nuclei, cytoplasm-objects using the original (non preprocessed) fluorescent images of DAPI, FUS, SFPQ and ChAT.

#### Post-mortem issue (Supplementary Fig. 1B)

*Preprocessing:* to facilitate MN segmentation, we first enhanced the contrast and equalized the histogram of the fluorescent channels (DAPI, ChAT, FUS) using existing packages within FiJi (ImageJ - see preprocessing scripts provided in Zenodo), which served to counteract the varying background noise of the images. These preprocessing steps have been adopted to facilitate object detection, and the modified images were not used in the following automated analysis steps that directly measure staining intensity or densitometry.

*Pixel Classification:* A randomly selected representative subset of the preprocessed images (26 in total) was then used for training a parallel random forest (VIGRA) algorithm in Ilastik (see labels and training set examples provided in Zenodo) to identify MNs from background, and to exclude the corpora amylacea (age-related insoluble accumulations detectable in postmortem tissue that tend to present non-specific immunolabeling across all channels, and need to therefore be identified and excluded from the analysis, henceforth identified as “inclusions”). The training was performed on preprocessed images using 3-channels (DAPI, ChAT, FUS).

*Automated Image Analysis:* The binary masks generated by the pixel classification (ML-MNs) were then added to the original channels and used as a dataset for automated analysis in Cellprofiler (see pipeline provided in Zenodo). In Cellprofiler we implemented steps to detect nuclei as a secondary structure based on the MNs identified in Ilastik (ML-MNs) overlaid to the DAPI channels, and the cytoplasm was determined by subtraction of DAPI to the identified MNs as a tertiary structure. The resulting nuclei- and cytoplasm-objects were used to measure the texture, intensity, intensity distribution, size, shape, colocalization and radial distribution for each MN using the (non preprocessed) fluorescent images of DAPI, FUS, SFPQ and ChAT.

### Preprocessing of high-content imaging data

Measurements from whole MNs, nuclear and cytoplasmic compartments were merged to form a single data matrix. Subsequent analyses were performed with the R statistical package version 3.3.1, Bioconductor libraries version 3.3 (R Core Team. R: A Language and Environment for Statistical Computing. Vienna, Austria: R Foundation for Statistical Computing; 2013), and Python 3.5.3 in a single Jupyter notebook framework. MNs with cytoplasmic compartment areas smaller than nuclear compartment areas were removed for primary analysis in order to ensure similar positioning of the cells. As normal distributions make it easier to work with numeric values from a mathematical, statistical, and computational point of view(Caicedo *et al*., 2017), we iteratively tested the normality of the measurements across the cells and log-transformed to obtain approximate normal distributions for features that have highly skewed values or require range correction. Additionally we standardized the measurements to ensure approximately normally distributed, mean centered and have standard-deviation.

### Unsupervised characterisation of high-content microscopy data

Hierarchical clustering of the cells based on their morphological profiles has been carried out using weighted-average linkage applied to euclidean distances. Singular Value Decomposition (SVD) has been performed on the hundreds of measurements across hundreds of cells. Selection of the components maximally capturing variance in gene expression resulted in a subset of components to focus on. Linear mixed model (LMM) was then used to test the association between each of the *n* first selected principal components and vcpALS phenotype (VCP-mutant MNs), comALS phenotype (VCP- and SOD1-mutant MNs), or sALS phenotype, accounting for idiosyncratic variations due to the animals or individuals where the cells originated. The right singular vectors were used to generate the PCA scatter plots of the MNs projected on the principal components, while the left singular vectors were used to extract the contribution of each measurement to each component.

### Automated identification of MNs subpopulation

Gaussian Mixture Models are a powerful label-free probabilistic distribution-based clustering algorithm that have been shown to be effective in capturing subpopulations in imaging data(Slack *et al*., 2008, Loo *et al*., 2009*b*). In this model, the subpopulations and their proportions correspond to mixture centers and mixture prior probabilities, respectively. Both of these quantities are considered as unknown parameters and were estimated using the expectation maximization (EM) algorithm. The algorithm was initialized with unit covariance and the centroid positions obtained using the k-means algorithm. The starting positions of the centroids in the k-means algorithm were initialized randomly, meaning the algorithm is nondeterministic.

To compare against the label-independent GMM-based classifier, we also selected two label-dependent probabilistic classifiers, namely Logistic Regression (LR) and a multilayer perceptron (MLP) neural network with one hidden layer. LR and MLP can deal with more complex patterns in data whilst relying on labeling of the cells. In particular MLPs decision boundaries can be nonlinear. In LR the contribution of parameters (coefficients and intercept) can be easily interpreted, which is not always the case with the parameters of a neural network. Prior to training, we splitted the data into train (70% of the cells) and test sets (30% of the cells) in stratified fashion by conserving the relative fraction of cells originating from each animal or individual. The regularization strength of the LR has been optimised using 10-fold cross-validation over a parameter grid and selecting against the best accuracy score. The hyperparameters of the MLP, namely the regularization term, the hidden layer size, the solver and the activation were selected using 5-fold cross-validation over a parameter space and selecting against the best accuracy score. GMM, LR and MLP have been trained and optimised in the SciPy environment using the scikit-learn library (Pedregosa *et al*., 2011). The performances of each classifier were then compared using the Receiver Operational Characteristics (ROC) curves and the Area under the curves (AUC).

### Relative contribution of cell measurements

The relative contributions of the individual measurements to the different classifiers were obtained as follows. For GMM-based classifiers, the left singular vectors of the components used for GMM modeling were extracted and then converted in relative quantities by dividing each vector by its sum. For LR classifiers, the weights were extracted and similarly converted into relative quantities by dividing each weight by the sum of all the weights for a given LR. MLP uses equation (1) to model the probability of each MNs to be sick given the observed measurement vector X of dimension [1 × *m*] where *m* is the number of measurements, *w_i_* is the weight of the *i*th perceptron of the hidden layer, *v_ij_* is the weight of the *j*th cell measurement in the *i*th hidden perceptron, and *n* is the number of perceptrons in the hidden layer.

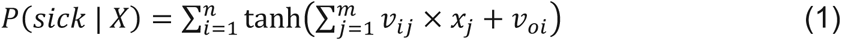

To calculate the relative contribution 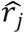 of each measurement *j* to the MLP classifier, we first extracted the global relative contribution using equation (2).

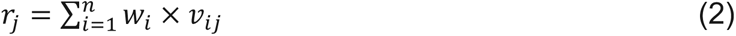

In order to get then the relative contribution to the classifier, we finally divided all contributions by the sum of all contributors as shown in equation (3).

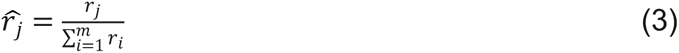

### Scoring metrics

GMM, LR and MLP are probabilistic classifiers which output a per-cell posterior probability *P* to be sick given the observed phenotype. The biological interpretation of this probability relates to the confidence for a given cell to be either sick or healthy given the model. We also transformed these disease probabilities using equation (4) to generate alternative scoring metrics that relate to the ‘disease severity’ expected to better reflect on the possibility for two cells to exhibit very different degrees of aberrant phenotypes while having similarly high probability to be sick (**Supplementary Fig. 3**).

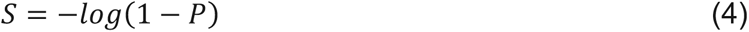

The per-cell disease profiles were finally constructed using 1) cell-level probability of the disease and 2) cell-level disease severity score. These vectors are expected to capture the disease status of each cell as captured by the cellular morphology. From these two metrics we also extracted 1) per-animal probability to be sick by averaging the probabilities across all cells for each animal, 2) per-animal disease severity by averaging the severity scores across all cells for each animal, 3) per-animal fraction of sick cells by computing the fraction of cells with probability above 0.0 to be sick in each animal.

### Data availability

We provide raw images and complete source code (which is not a software but rather a compilation of R and python) to readily reproduce figures, tables, and other results that involve computation in order to facilitate the development and evaluation of additional profiling methods. We also provide the measurements of each of the ∼600 cells whose origins are annotated. The raw images, metadata and single-cell measurements provided as comma-delimited files have been deposited <u>Zenodo</u> under the accession number 3985099, together with the image processing pipelines. The scripts for automated detection of MNS subpopulation can be freely accessed on Github in the following repository: https://github.com/RLuisier/ALSdisMNs.

## AUTHOR CONTRIBUTIONS

Conceptualization, R.L., A.S., R.P.; Formal Analysis, C.H., A.S., R.L.; Investigation, C.H., G.E.T., R.L., A.S.; Writing – Original Draft, R.L., A.S., R.P.; Writing – Review & Editing, R.L., A.S., R.P., C.H., G.E.T., J.N., H.D.; Human sample provision, J.N.; Resources, R.P.; Visualization, R.L.; Funding Acquisition, R.L., A.S., R.P.; Supervision, R.L., A.S., R.P.

## ACKNOWLEDGMENTS

This work was supported by the Idiap Research Institute and by the Francis Crick Institute which receives its core funding from Cancer Research UK (FC010110), the UK Medical Research Council (FC010110), and the Wellcome Trust (FC010110). R.P. holds an MRC Senior Clinical Fellowship [MR/S006591/1]. A.S acknowledges the support of the Wellcome Trust [213949/Z/18/Z].

